# Spatiotemporal Proteomics Deciphers Functional Selectivity of EGFR ligands

**DOI:** 10.1101/2023.09.13.557663

**Authors:** Akihiro Eguchi, Kristina B. Emdal, Zilu Ye, Ulises H. Guzmán, Jesper V. Olsen

## Abstract

The epidermal growth factor receptor (EGFR) induces different signaling outputs depending on ligand identity and biological context. This phenomenon is known as functional selectivity, but the underlying molecular mechanisms remain elusive. Here, we investigated this on a global scale and time-resolved by high-throughput multilayered proteomics integrating dynamic changes in the EGFR interaction network by proximity biotinylation using EGFR- TurboID, phosphoproteome, and proteome in response to stimulation with the six highest-affinity EGFR ligands. We obtained comprehensive temporal profiles of protein recruitment and phosphosite changes pinpointing signaling proteins differentially regulated by the six ligands with key impact on EGFR endocytic fate, e.g. degradation or recycling. Specifically, the Epsin family protein, Clint1 was identified to control the endocytic trafficking of EGFR towards degradation. Moreover, we characterized the protein interaction selectivity of EGFR C-terminally phosphorylated tyrosine residues using a panel of tyrosine mutated constructs showing STAT5 specificity for EGFR Y1173. These data provide a comprehensive resource deciphering functional selectivity of EGFR signaling to support discovery of novel drug targets.

## Introduction

Functional selectivity is a phenomenon defined as the ligand-dependent selectivity for particular cell signaling pathways when ligands activate the same receptor, and the degree to which each pathway is activated depends on which ligand binds to the receptor^1^. The term was first established for G-protein coupled receptors, for which functionally selective ligands were initially reported, and later cytokine receptors as well as receptor tyrosine kinases (RTKs) such as c-Met^2^, epidermal growth factor receptor (EGFR)^3-6^, and fibroblast growth factor receptor (FGFR)^7^ were identified to also have several ligands with biased signaling abilities. In drug discovery, understanding the molecular mechanisms underlying functional selectivity holds great impact, because this mechanism raises the possibility of selecting or designing novel ligands that differentially activate and eventually control the “strength of signaling” for only a subset of functions of a single receptor. Thus, enabling more optimal, beneficial, and fine-tuned therapeutic responses. Promising preclinical examples of functionally biased ligands are the insulin analog S597 showing differential activation of metabolic signaling pathways downstream of the insulin receptor^8^ and the engineered interleukin-2 inducing proliferation of CD8-positive T cells without differentiating them into effector T cells^9^.

Among RTKs, EGFR serves as the prime receptor system for studying functional selectivity given its 7 known ligands and their ability to elicit differential cellular signaling outcomes. EGFR is expressed in various cells and tissues and transduces diverse signaling depending on the biological context. Critically, numerous cancers display EGFR overexpression or mutations with oncogenic autonomous signaling abilities rendering EGFR a prominent oncoprotein and tractable target for anti-cancer therapy^6, 10–12^. However, canonical activation of EGFR is ligand-dependent, and upon binding of the classical ligand EGF, the receptor dimerizes to transphosphorylate intracellular kinase domains, and ultimately, phosphorylated tyrosine residues in EGFR function as docking sites of the intracellular signaling proteins that activate downstream signaling. To date, seven ligands for EGFR have been identified (EGF, betacellulin (BTC), heparin-binding EGF (HB-EGF), transforming growth factor α (TGFα), amphiregulin (AR), epiregulin (ER), and epigen (EPI)). Upon activation, EGFR is internalized to early endosomes, and sorted to either recycling endosomes destined to reappear on the cell surface or late endosomes that fuse with lysosomes leading to lysosomal degradation of EGFR^13^. Endocytosis of EGFR occurs via a wide variety of trafficking pathways determining the endocytic EGFR fate. These can be distinguished by recruitment of different sorting protein machineries involving adaptor proteins such as clathrin, AP-2, and CBL^14–17^. The balance of receptor recycling and degradation depends on the type of extracellular ligands, and Roepstroff et al. categorized six EGFR ligands with the highest affinity to EGFR as degradation-inducing ligands (hereafter DEG-ligands; EGF, BTC, and HB-EGF) and recycling-inducing ligands (hereafter REC-ligands; TGFα, AR, and ER)^18^. Consequently, receptor trafficking involves a complex network of protein-protein interactions, post-translational modifications including phosphorylation, and it is the coordinated recruitment of proteins to EGFR that ultimately determine receptor fate. Therefore, a spatiotemporal resolution during the course of receptor trafficking is required to understand the molecular mechanisms underlying functional selectivity in the context of EGFR.

A variety of mechanisms may influence functional selectivity including ligand identity and receptor binding abilities, differences in ligand-induced conformational states, receptor homo/hetero dimerization, and the recruitment profile of proximal scaffolding proteins and activation of downstream signaling proteins^5, 6^. However, reported studies aimed at deciphering the molecular events underlying EGFR functional selectivity have largely focused on a single or at best two ligands, and with limited spatial information^19-22^. Francavilla et al. performed a multi-layered proteomics analysis comparing signaling responses induced by EGF and TGFα, and identified phosphorylation of late endosomal marker protein Rab7 and recruitment of recycling endosomal marker proteins Rab11Fip1 as key for the degradation and recycling process, respectively^19^. Verdaguel et al. characterized spatiotemporal profiles of proximal proteome in response to EGF and showed the Trk-fused gene (TFG) is a regulator of endosomal sorting of EGFR^21^. Knowing the importance of the tight regulation of intracellular signaling networks for cellular decision-making underscores the necessity to study the specific timing of well-coordinated molecular events. Therefore, systematic profiling of all EGFR ligands on a proteome-wide scale with spatiotemporal resolution is needed to uncover the molecular complexity of EGFR functional selectivity.

Here, we defined and enabled an EGFR-centric approach to create a comprehensive resource of three signaling layers describing functional selectivity. This included dynamic mapping of interaction networks, phosphoproteomes and proteomes in response to the six highest-affinity ligands in a cell system supporting differential EGFR signaling and fates. The preferred method for spatiotemporal interaction network mapping utilizes a biotin ligase-based proximity labeling technique^23–26^ and therefore, we fused a biotin ligase enzyme TurboID^24^ to the C-terminus of EGFR to allow for biotinylation of recruited proteins in close proximity and identification by affinity-purification (AP)-mass spectrometry (MS). Using this approach, we temporally tracked cellular signaling events induced by six natural EGFR ligands and linked this information to the endocytic fate of the receptor. Specifically, we identified a role for the clathrin interactor protein CLINT1 as a molecular switch determining the endocytic receptor fate. Moreover, we mapped the selectivity of tyrosine residues in the EGFR C-terminal tail using tyrosine-mutated EGFR-TurboID constructs. Besides covering a large part of the known phosphotyrosine-binding protein network, we revealed the specific binding of STAT5 to EGFR pY1173. Our approach provides a vast resource of molecular networks of EGFR signaling for a deeper understanding of the molecular mechanisms underlying functional selectivity.

## Results

### Development and validation of the EGFR-TurboID system to study EGFR trafficking

To address functional selectivity in the context of EGFR signaling and its natural high-affinity ligands, we set out to characterize comprehensive time-resolved ligand responses at the level of PPI networks, phosphoproteome and proteome. Time points for ligand stimulation were chosen to cover signaling events from initial EGFR activation to receptor endocytosis through differential trafficking routes e.g. degradation and recycling (Fig. 1A). The HeLa cervix carcinoma cell line is a universal model for studying EGFR signaling at physiological receptor levels, and ligand-induced EGFR trafficking has mainly been characterized in these cells. Consequently, to study EGFR proximity labeling-based interaction networks analysis using MS-based proteomics, we initially established a HeLa cell clone stably expressing an engineered EGFR-TurboID fusion construct. Overexpression of a promiscuous biotin ligase often causes cell toxicity due to excessive and accumulated protein biotinylation. Thus, we used a tetracycline (TET)-inducible expression system to control the EGFR-TurboID expression level^27, 28^. We transfected HeLa Flp-in TREX cells with a plasmid encoding an EGFR-TurboID construct and established a cell clone named T1 by limiting dilution. We confirmed by western blotting that clone T1 showed a time-dependent increase in expression level of EGFR-TurboID upon TET-induction (Fig. 1B). In agreement with previous reports^29^, TET-inducible expression systems can be challenged by leaky expression and accordingly, our clone showed leaky expression of EGFR-TurboID in the absence of TET at near-endogenous receptor levels (Fig. 1B). Only the leaky state and not the TET-induced conditions, allowed for the concomitant degradation of both endogenous EGFR and the EGFR-TurboID fusion construct upon treatment with EGF and BTC (Fig. 1C). Consequently, we speculated that the TET-induced levels of EGFR-TurboID exceeded the capacity of the cellular EGFR trafficking machinery, underscoring the importance of a near endogenous level of expression for the fusion construct to properly mirror endogenous EGFR biology. Given that clone T1 meets this criteria, we deemed this clone a good candidate for further characterization and thus, compared the dose-response effects of short-term EGF treatment in clone T1 and HeLa cells. Reassuringly, we confirmed not only comparable EGFR phosphorylation status saturating at approximately 5–10 nM EGF but also similar activation of downstream signaling proteins Akt and Erk1/2 by western blotting (Fig. S1A). To fully assess the ability of the EGFR-TurboID fusion to activate downstream signaling in absence of endogenous EGFR, we transiently expressed the fusion construct in A549 EGFR-knockout cells and confirmed activation of Erk1/2 upon EGF stimulation (Fig. S1B). Next, we evaluated EGFR internalization and confirmed by immunostaining and confocal microscopy EGFR presence at the cell surface and co-localization in distinct intracellular punctae with the early endosome marker protein EEA1^30^ upon 8 min EGF stimulation in clone T1 (Fig. 1D). Moreover, EGFR-TurboID showed the same pattern when staining for TurboID and biotinylation upon EGF stimulation (Fig. 1E), thus, confirming the ability of the fusion to undergo internalization under active biotinylation conditions. These results addressing kinase activity, downstream signaling, EGFR degradation and internalization in response to ligand, show minimal deviation from the endogenous EGFR response. Therefore, we deemed clone T1 with EGFR-TurboID expression at endogenous levels a suitable cell model for subsequent large-scale proteomics analyses addressing questions related to EGFR signaling and differential trafficking responses of the ligand repertoire.

**Figure 1.**
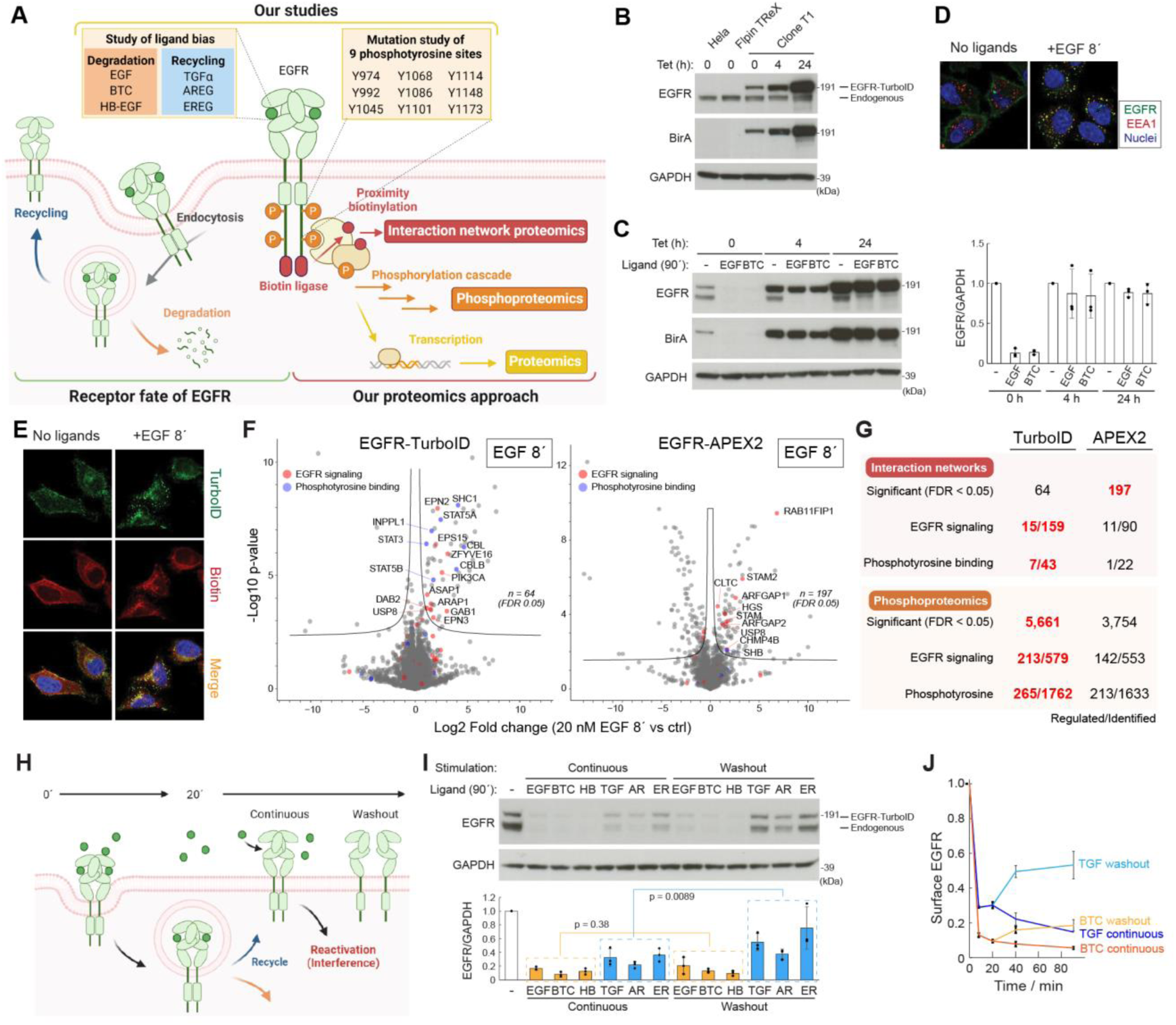
(A) Multi-layered proteomics approach for the comprehensive dissection of EGFR signaling pathway. (B) Western blotting of the cell lysates of HeLa, HeLa-FlpIn, and clone T1 cells treated with TET for indicated time. (C) Western blotting of the cell lysates of clone T1 cells stimulated with EGF (20 nM) or BTC (20 nM) for 90 min after the TET induction for indicated time. Values are mean ± s.d. (n = 3). (D) Confocal microscopic images showing EGFR internalization. (E) Confocal microscopic images showing colocalization of EGFR and biotinylated proteins. (F) Volcano plots highlighting EGFR signaling-related proteins (red), KEGG term: “ErbB Signaling Pathway” and/or “Endocytosis”) and proteins with phosphotyrosine binding domains (blue, Pfam term: “Cbl_N2”, “SH2”, and/or “PTB”) from interaction network analysis data using EGFR-TurboID- or EGFR-APEX2-expressing cells. Fold change represents EGF treatment (20 nM, 8’) versus control. (G) Summary of the numbers of regulated or identified proteins/phosphopeptides. (H) Illustration of the interference by remaining ligands and our stimulation methods. (I) Western blotting of the cell lysates of T1 cells stimulated with indicated ligands for 90 min using continuous or washout method. Values are mean ± s.d. (n = 3). (J) Flow cytometry analysis showing the surface expression level of EGFR after the treatment with BTC (20 nM) or TGFα (20 nM). Values are mean ± s.d (n = 3).

### Comparative analysis of EGFR-TurboID and EGFR-APEX show superior identification of known EGFR signaling-related proteins/phosphopeptides for EGFR-TurboID

For protein interaction studies using proximity labeling, two major types of biotinylating enzymes exist, the biotin ligases (e.g. BioID and TurboID)^23, 24^ and ascorbate peroxidase (APEX and APEX2)^25, 26^. Biotin ligases use biotin as a substrate and mainly label lysine residues in target proteins with reaction time in minutes to hours, whereas ascorbate peroxidases use biotin phenol to label tyrosine residues within a minute after the supplement of hydrogen peroxide as an initiator. Here, we initially evaluated ligand-dependent proximity-labeling of known EGFR interactions by fusing EGFR to either of the two biotinylating enzymes, EGFR-TurboID and EGFR-APEX2. We established cell clones with TET-inducible expression of EGFR-APEX2 as described for EGFR-TurboID, and similarly selected a clone A1 with an EGFR-APEX2 expression level comparable to endogenous EGFR, however, requiring a 6 h TET-induction (Fig. S1C). The ability of the two constructs to biotinylate proteins *in vivo* was confirmed by western blotting, and the addition of biotin (TurboID) or biotin-phenol and H_2_O_2_ (APEX2) was required for increased biotinylation levels (Fig. S1D).

For comparative interaction proteomics and phosphoproteomics, we prepared cell lysates after 8 min treatment of EGF in presence or absence of the EGFR kinase inhibitor erlotinib, and processed samples following the workflows as described in Fig. S1E. We evaluated significantly regulated proximal proteins and phosphorylation levels upon EGF stimulation by t-test statistics visualized by volcano plots and identified 64 and 197 proteins and 5,661 and 3,754 significantly regulated phosphopeptides for EGFR-TurboID and EGFR-APEX2, respectively (Fig. 1E and Fig. S1F). Although LC-MS/MS analysis of biotinylated proteins enriched from both Turbo-ID and APEX2 expressing cells identified thousands of potentially proximal proteins, applying a quantitative filter based on fold-change significance effectively prioritizes true interactors over background binders. A greater number of EGF-dependent proximal proteins showed a known association with EGFR signaling and contained phosphotyrosine-binding domains for EGFR-TurboID compared to EGFR-APEX2 (Fig. 1F and 1G). For the phosphoproteome, samples derived from EGFR-TurboID showed an overall greater number of identified phosphorylated peptides also reflected in phosphorylated peptides derived from EGFR signaling proteins and containing tyrosine phosphorylation (Fig. 1G and Fig. S1F). Overall, these results supported our choice of EGFR-TurboID for large-scale proteomics with the added advantage of the simplest biotinylation stimulation and cell lysis workflow compared to EGFR-APEX2. A simple setup favoring a great coverage of EGFR-related proteins support scalability and quality of our proposed analyses for studying EGFR biology.

### Six highest-affinity ligands of EGFR induce different receptor fates: degradation or recycling for EGFR-TurboID expression

To elucidate the molecular mechanisms of functional selectivity, we evaluated the effect of the six highest-affinity ligands of EGFR: EGF, BTC, HB-EGF, TGFα, AR, and ER in clone T1. We excluded EPI due to the fact that it required a very high concentration (> 1 µM) to activate EGFR in our system (data not shown). Firstly, to identify the individual ligand concentrations allowing for comparable EGFR activation levels, we treated T1 cells with a concentration range of each ligand for 5 min and assessed the degree of phosphorylation of EGFR and downstream signaling by western blotting (Fig. S1G). Based on these results, we decided to use 20 nM for BTC, TGFα, and HB-EGF and 500 nM for AR and ER for which phosphorylation of EGFR reached a maximum and saturation. Secondly, we aimed to confirm the categorization (degradation versus recycling) of the ligands at these ligand-specific concentrations, and monitored the magnitude of EGFR degradation and surface expression after up to 90 min stimulation by western blotting and flow cytometry, respectively (Fig. 1H–J). EGFR is known to recycle to the cell surface after 20–30 min of ligand stimulation^18^, and REC-ligands remaining in the culture medium can reactivate the recycled receptors within 90 min stimulation. Due to this reactivation, we cannot distinguish and compare first-wave EGFR activation to that of DEG-ligands. Thus, to better define this state for REC-ligands, we compared two stimulation methods: one with “continuous” ligand presence and a “washout” condition removing ligands after 20 min stimulation (Fig 1H). We found that the three DEG-ligands (EGF, BTC and HB-EGF) to a greater extent reduce EGFR levels (endogenous and fusion) in the T1 clone compared to the REC-ligands (AR, ER and TGF), and this was even more pronounced for the washout condition indicating a greater degree of recycling upon first-wave activation only (Fig. 1I). These findings were confirmed by EGFR cell surface expression for BTC and TGF (Fig.1J) and validated that in the T1 clone, differential EGFR endocytic fates in response to the different EGFR ligands is observed and thus, establish our model as well-suited for our large-scale proteomics aiming to address functional selectivity.

### Multilayered proteomics analysis identifies temporal changes induced by EGFR activation

Next, we prepared samples for multilayered large-scale proteomics using the T1 cell clone. The experimental setup included a total of 300 samples consisting of lysates from cells after stimulation with one of the six ligands for 3, 8, 20, 30, 40, or 90 min (Fig. 2A). The 40 min and 90 min stimulation included continuous and washout conditions. As controls, we included a no biotin condition, a no-ligand-stimulated condition, and a condition of 8 min ligand stimulation after 15 min pre-incubation with erlotinib. Each condition was performed in five biological replicates and biotin was added to the culture medium 10 min prior to harvest for all conditions except the no biotin control. The sample processing workflows for our high-throughput large-scale proteomics (interaction network, phosphoproteome and proteome analysis) were semi-automated utilizing a 96-well plate format and the KingFisher robot prior to LC-MS/MS analysis. Cell lysate from each of 300 samples was divided into three different plates and processed as illustrated (Fig. 2A), according to the individual proteomics sample preparation workflows. In brief, the proximity network proteome analysis consisted of streptavidin (SA) bead-based enrichment of biotinylated proteins followed by on-bead Lys-C/trypsin protein digest. The phosphoproteome analysis included protein aggregation capture (PAC)^31^ and on-bead Lys-C/trypsin protein digestion followed by phosphopeptide enrichment by immobilized metal affinity chromatography using titanium ions (Ti-IMAC), and the proteome analysis was conducted after on-bead protein digest. All resulting peptide mixtures were analyzed by LC-MS/MS with the Evosep One LC coupled to an Orbitrap Exploris 480 mass spectrometer operated in data-independent acquisition (DIA)-mode utilizing 30 samples per day (SPD) (interaction proteome and proteome) or 60 SPD (phosphoproteome) LC gradients.

**Figure 2.**
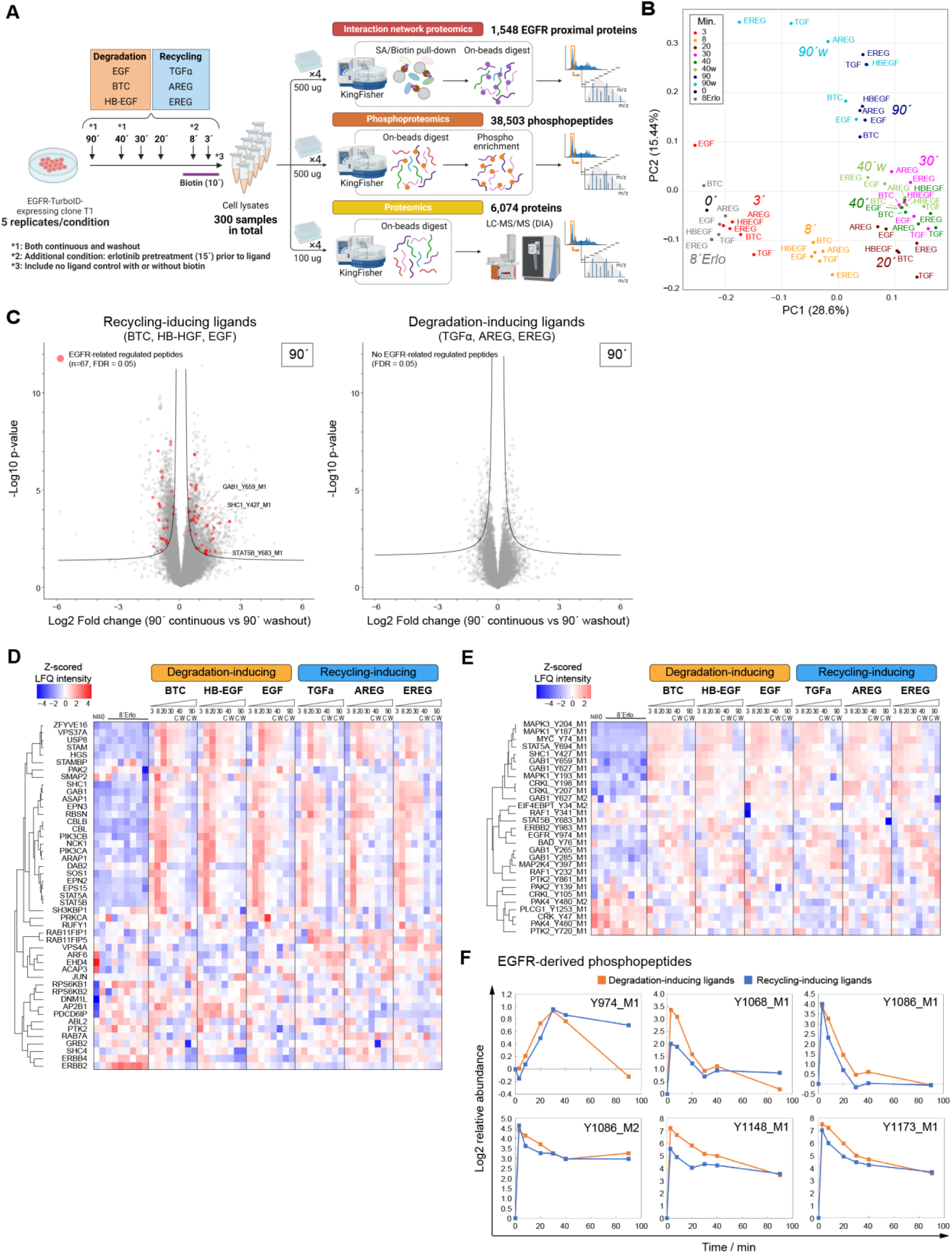
(A) Schematic illustration of the multilayered proteomics approach. (B) Principal Component Analysis (PCA) of phosphoproteomics data acquired at various time points. (C) Volcano plots showing differentially regulated phosphopeptides for each ligand group. Fold change represents 90 min stimulation with washout method versus continuous method. (D, E) Heatmaps of EGFR signaling-related proteins/phosphopeptides from (D) interaction network analysis data and (E) phosphoproteomics data. (F) Temporal profiles of log2 abundance of EGFR-derived phosphopeptides. The values are subtracted by the values at 0 min.

After data filtering, we identified and quantified 1,548 EGFR proximal proteins from the interaction network analysis, 38,503 phosphopeptides (class I^32^, 4% pTyr: 1,535, 96% pS/T: 36,968) from the phosphoproteome, and 6,074 proteins from the proteome (Fig. 2A and Fig. S2A). Principal component analysis (PCA) of the datasets revealed a pronounced data clustering according to ligand grouping (DEG versus REC) for the interaction analysis (Fig. S2B), whereas phosphoproteome data clustered according to time of stimulation irrespective of ligand identity (Fig. 2B). This suggests that proximal proteins rather than global phosphoproteome dynamics hold promising potential to differentiate DEG- from REC-ligands. Noteworthy, ligand washout conditions clearly separated from the continuous stimulation, particularly in the phosphoproteome data, and REC-ligands showed separation that is more pronounced at the 90 min time point compared to DEG-ligands (Fig. 2B and Fig. S2B). Volcano plots showing upregulation of numbers of proximal proteins confirmed this observation (96 versus 38; FDR < 0.05, Fig. S2C) and phosphopeptides (67 versus none; FDR < 0.05, Fig. 2C) among proteins related to EGFR signaling when comparing the 90 min time point versus unstimulated control for REC- and DEG-ligands. These observations matched expectations given that the washout condition removes the ligand, thus, eliminating reactivation of recycled receptors at the cell surface.

With this data, we could detect and confirm regulation of EGFR signaling-related proximal proteins and their phosphorylation as exemplified by volcano plot analysis showing fold change differences and significance comparing the 20 min EGF stimulation and non-stimulated control. (Fig. S2D). To analyze the temporal profiles of proximal proteins and phosphotyrosine-containing peptides related to EGFR signaling, we created heatmaps with z-scored median log2-transformed LFQ intensities (Fig. 2D and Fig. 2E). This confirmed the ability to map proximal dynamics of many known direct EGFR interactors. For example, CBL, SOS, STATs, and EPNs were predominantly/maximally recruited at early time points around 3–8 min, and proteins belonging to the endosomal sorting complexes required for transport (ESCRT) such as HGS, STAM, and USP8 peaked at later time points around 20–40 min (Fig. 2D) ^33^. These results confirmed that our interaction network analysis provided informative temporal profiles for known EGFR signaling proteins and proteins involved in receptor trafficking. Noteworthy, we confirmed the specific recruitment of RAB11FIP1 to EGFR in response to all REC-ligands and not DEG-ligands. We have previously reported Rab11Fip1 to serve as a key protein for transferring EGFR to recycling endosomes in response to TGFα^19^. In the phosphoproteome data, known tyrosine phosphorylation sites of STAT5, MAPK, and SHC1 showed similar transient activation profiles (Fig. 2E). Among nine potential phosphotyrosine sites of EGFR, five phosphosites from six peptides were successfully quantified (Fig. 2F). Their temporal profiles were early transient except for pY974 showing a late transient profile. In general, DEG-ligands induced a higher level of phosphorylation than REC-ligands, this was especially prominent for the pY1068 site, which we confirmed by western blot analysis (Fig. S2A). Taken together, we have highlighted and confirmed temporally resolved known EGFR signaling and biology for each of three proteomics layers. As a consequence of a biologically fine-tuned and validated model system, we deemed the data of high quality and suitable for subsequent in-depth characterization and hypothesis-generation.

### Interaction network analysis reveals ligand-specific dynamic protein profiles and proximal proteins related to clathrin-coated vesicles with important roles in EGFR recycling

To focus on proximal proteins differently regulated between REC- and DEG-ligands, we initially calculated two main averaged profiles for each proximal protein by averaging their log2-transformed LFQ intensities for each time point for each group of REC- and DEG-ligands, respectively. Next, we performed fuzzy c-means clustering based on the temporal profiles from 909 proteins with strong upregulation (>1.5 fold change) in the interaction network data (Fig. S3A), which resulted in eight distinct clusters of temporal profiles (Fig. 3A). Venn diagrams showed unique and overlapping degradation and recycling profiles for each cluster (Fig. 3A). Moreover, we summarized the overlap of members within each cluster in the Sankey diagram in Fig. 3B. From this, it was evident that some proximal proteins belong to different temporal clusters dependent on degradative or recycling ligand stimuli, whereas a large part of proximal proteins is assigned to the same cluster. Hence, with this data, we are able to quantify and track proximal protein dynamics specific to either ligand group. Enrichment analysis of Gene Ontology (GO) and Kyoto Encyclopedia of Genes and Genomes (KEGG) pathway terms for each cluster revealed overrepresentation of cytoplasmic proteins mainly in clusters 1 to 6, and nuclear proteins in clusters 7 and 8 (Fig. 3C and S3B). In particular, cluster 1, with early transient signaling (peak at 8 min ligand stimulation), showed the most pronounced enrichment of proteins related to ErbB signaling pathway, endocytosis, and clathrin-coated vesicles (CCVs), whereas cluster 2, with late transient signaling (peak at 20 min ligand stimulation), showed predominant enrichment of proteins related with endosomes. These findings align well with proteins in cluster 1 being involved in EGFR activation and internalization, whereas cluster 2 is enriched for proteins involved in timing of EGFR trafficking and fate. Clusters 3 to 4 showed no significant enrichment of terms, whereas clusters 5 and 6 were enriched in chromatin-related proteins and cellular junction-related proteins, respectively. Noteworthy, the largest overlap in protein DEG- and REC-profiles was seen for nuclear proteins in clusters 7 and 8. (Fig. 3B and S3B). We were surprised to see the dynamics of this class of proteins in the interaction data, however, a profile comparison to protein abundance profiles from the proteome data revealed a striking profile similarity (Fig. S3C). Hence, we speculate that this group of proteins are nucleo-cytoplasmic shuttling. We applied a cell lysis protocol with modified RIPA to favor the cytoplasmic compartment for our big setup and thus, this serves a reasonable explanation for our cluster 7 and 8 findings.

**Figure 3.**
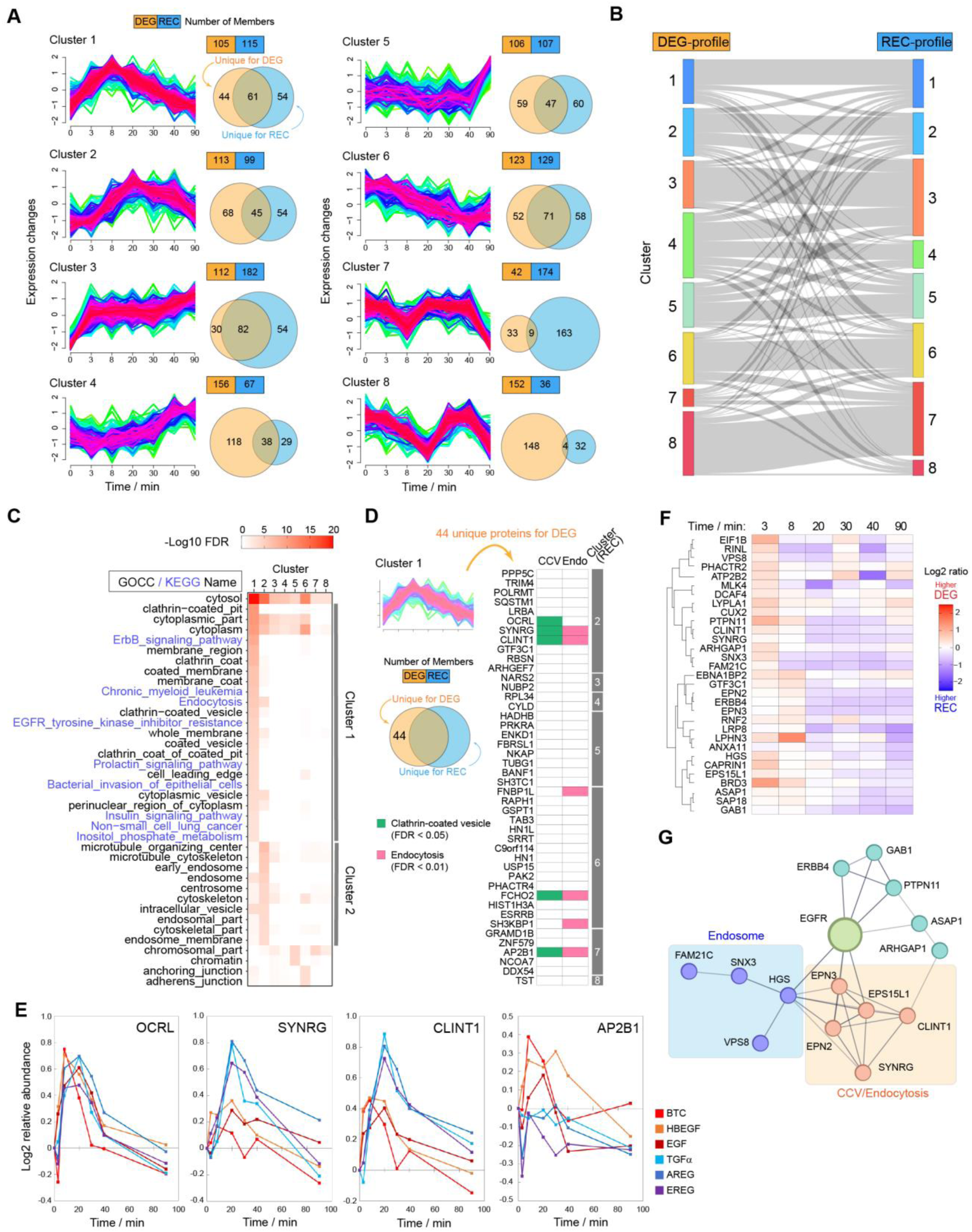
(A) Fuzzy c-means clustering of EGFR proximal proteins. Each protein has two lines representing the respective profiles induced by the degradation- and recycling-inducing ligands. The numbers of members for each cluster are shown in the Venn diagrams. (B) Sankey diagram describing the association of clusters between degradation- and recycling-inducing ligands. (C) GOBP and KEGG enrichment analysis based on the proteins in each cluster. See also Fig. S3C. (D) The list of 44 proteins assigned to cluster 1 only for degradation-inducing ligands. (E) Temporal profiles of recruitment of the proteins picked up from Fig. 3D. The values are standardized by the value at 0 min. (F) Clipped heatmap of the proteins differently regulated by two types of ligands. The color scale represents the difference between recycling-(blue) and degradation-(red) inducing ligands. See Fig. S3E for the whole heatmap. (G) Interaction network of the proteins listed in Fig. 3E. Only proteins that have at least one connection with other proteins are included. The network is created using the STRING database.

Based on the ligand-dependent interaction network dynamics observed, we hypothesized that proximal proteins with differential cluster assignments to be promising candidates as key players in functional selectivity. Given the enrichment of “ErbB signaling pathway” proteins in cluster 1, we analyzed the 44 proteins unique to DEG-ligands by GO enrichment analysis that revealed overrepresentation of proteins related to CCVs and endocytosis (Fig. 3D). Three of these differentially regulated EGFR proximal proteins, inositol polyphosphate 5-phosphatase OCRL (OCRL), clathrin interactor 1 (CLINT1), and synergin gamma (SYNRG) showed an early transient proximal cluster 1 profile for DEG-ligands and a late transient proximal cluster 2 profile for REC-ligands (Fig. 3D) and their individual ligand-specific profiles supported this finding (Fig. 3E). CLINT1 is an endocytic adaptor protein known to interact with the AP2 adaptor complex involved in the internalization of cargo in clathrin-mediated endocytosis^34, 35^, and we identified the beta subunit of AP2 (AP2B1) of this complex with an apex after 3 min in cluster 1 profile favoring recruitment to EGFR for only DEC-ligands (Fig. 3E). Hence, when applying a guilt-by-association principle, this data suggests that CLINT1, SYNRG, and AP2 are likely to be part of a complex specifically recruited to close proximity of EGFR when the receptor is activated by DEG-ligands. To define additional key proximal proteins specific to DEG- and REC-ligands, we calculated the log2-difference in LFQ intensities across the two ligand groups at each time point and visualized the candidate list of 243 proximal proteins with ratios favoring either of the two ligand groups in a heatmap (Fig. S3D). Cluster A represented proteins more strongly recruited by DEG-ligands at early time points (3 and 8 min) and by REC-ligands at later time points (20–90 min). The STRING network analysis of these proteins included not only CLINT1 and SYNRG but also additional clathrin-related endocytic proteins such as EPNs and EPS15 and endosomal proteins (Fig. 3F and 3G) ^35^.

### Clustering of regulated phosphopeptides reveals distinct and ligand-specific temporal profiles

To identify ligand-dependent phosphorylation site dynamics, we performed fuzzy c-means clustering of the regulated phosphoproteome data using the same approach as for the interaction network data analysis including two profiles, a DEG- and REG-profile for each phosphopeptide. Regulated phosphopeptides were assigned to 10 clusters, of which clusters 1–4 showed profiles with immediate early increases in phosphorylation levels and clusters 5–10 showed profiles with sub 20 min decline in phosphorylation levels (Fig. 4A and S4A). Mapping the ligand-dependent profile overlaps, we could confirm for the phosphoproteome data that all clusters showed some degree of shared profiles between DEG- and REC-ligands but also profile differences. Hence, dynamic differences in phosphoproteome profile may also reveal key signaling information addressing functional selectivity as shown for interaction data (Fig. S4B). With a focus on induced phosphorylation (clusters 1–4), we performed a kinase motif enrichment analysis using iceLogo to characterize the amino acid sequence (+/– 7) preferences surrounding the regulated phosphorylation sites within each ligand group (Fig. 4B and S4C). Cluster 1 with early and sustained phosphorylation from the 3 min time point, favored arginine in the –5 and –3 position, a consensus motif for kinases in the PKA, -B (Akt), and -C family^36^. Cluster 2 with transient phosphorylation around 8–40 min, favored proline at –2 and +1 position, a consensus motif for proline-directed kinases *e.g* MAPKs^36^. Cluster 3 with a delayed sustained profile showed ligand-specific motifs and despite a less pronounced enrichment for consensus motifs the DEG-ligands showed a mixture of PKA/B/C and proline-directed motifs which were absent for the REC-ligands (Fig. S4C). To infer the activity of immediate downstream kinases upon EGFR activation, we performed RoKAI (Robust Inference of Kinase Activity) analysis using profiles from cluster 1–4 at 3, 8, and 20 min after ligand stimulation (Fig. 4C). Ligand-dependent activation of different kinases was most evident for cluster 1 and 2 with several RTKs dominating in cluster 1 (green and purple in Fig. 4C). In accordance with the kinase motif enrichment analysis, many MAPKs are active in cluster 1 with ERK (MAPK1/3) and AKT1 showing stronger activation with REC-ligands. Conversely, cluster 3 only showed a few kinases with high activity, but encompasses the known stress response kinases, MAPK14 (p38), MAPKAPKs, and Akt2, which are activated predominantly by DEC-ligands (blue in Fig. 4C). These results suggest that phosphorylation of downstream proteins of MAPKs is either transient (cluster 2) or persistent (cluster 3), and that stress response kinases are persistently phosphorylated in the case of stimulation with degradation-inducing ligands. Immediate response phosphosites in Cluster 1 show more pronounced global kinase activity at 3 min compared to the other clusters. Compared to cluster 2, Akt1 and mTOR activities are particularly high, consistent with the kinase motif analyses (Fig. 4B).

**Figure 4.**
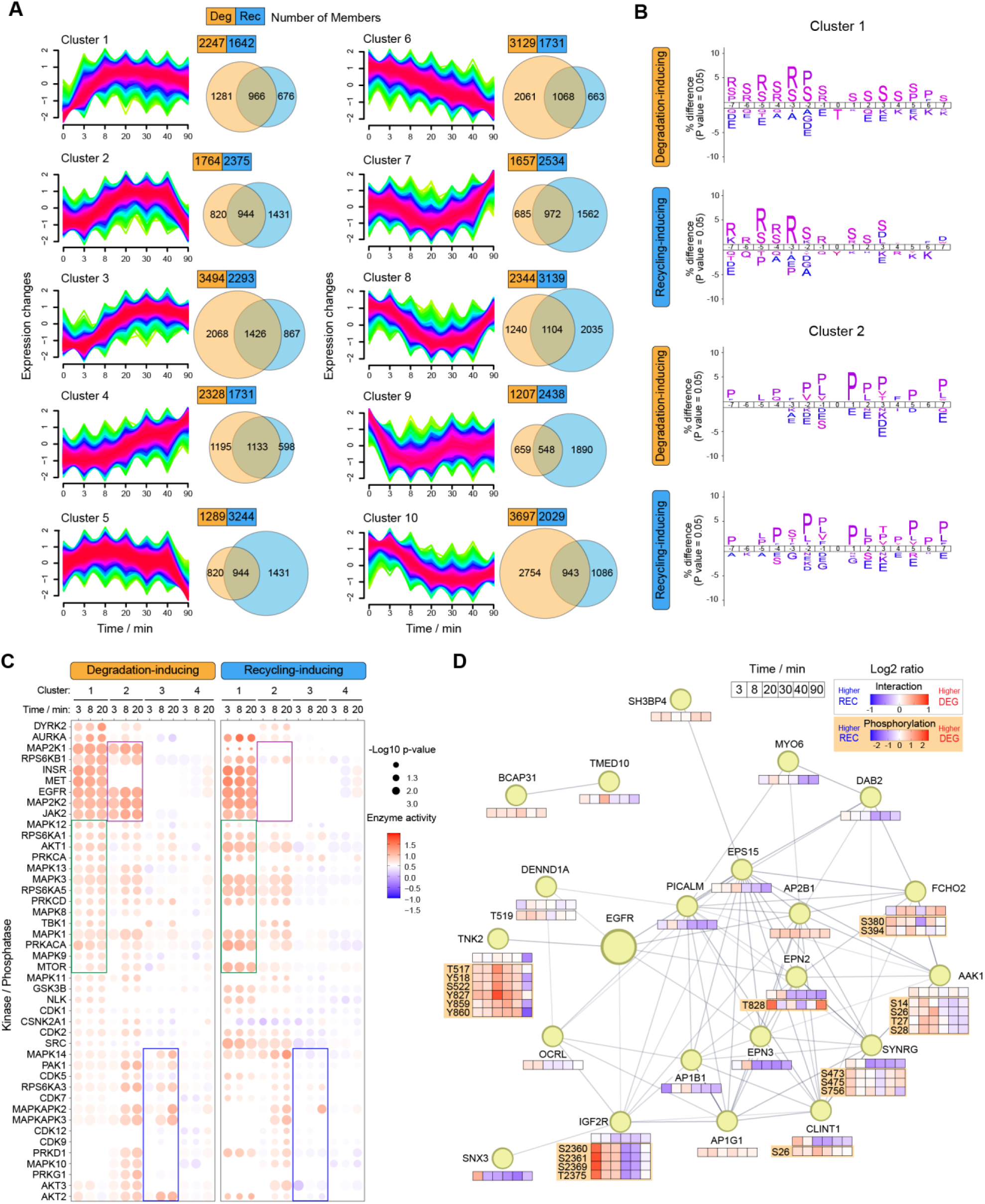
(A) Fuzzy c-means clustering of regulated phosphopeptides. Each peptide has two plots representing the respective profiles induced by the degradation- and recycling-inducing ligands. The numbers of members for each cluster are shown in the Venn diagrams. (B) Enriched phosphosites analysis using 15 sites surrounding phosphosites in clusters 1 and 2. The figures are created by IceLogo. (C) Kinase activity inference based on RoKAI using the abundance of phosphoproteins in clusters 1 to 4 for each time point. (D) Interaction network of CCV-related proteins with their profiles of recruitment and phosphorylation. The network is created using the STRING database.

To integrate interaction and phosphoproteome data, we focused on CCV-related proteins and created a functional association network based on STRING^37^. This analysis included regulated proximal EGFR proteins from the interaction data and phosphoproteins with phosphopeptides showing a >1.5-fold difference in their log2-transformed LFQ intensities between ligand groups for at least one time point (Fig.4D). Among key regulated phosphorylated sites showing a profile favoring DEG-ligands, we identified phosphorylation sites on the non-receptor tyrosine kinase, TNK2. We monitored two groups of two phosphotyrosine sites with opposing regulation patterns. The phosphorylated Y518 and Y827 sites displayed a decrease in levels upon ligand stimulation and REC-ligands induced this dephosphorylation earlier than DEG-ligands (Fig. S4D). In contrast, the known activating phosphorylation sites Y859 and Y860 showed induced levels by ligand stimulation, however, the duration of the phosphorylation was shorter for REC-ligands than DEG-ligands showing an earlier decline in levels. Intriguingly, the EGFR-TurboID recruitment profile of TNK2 was highly similar for all the ligands (Fig. S4E), which implies that phosphorylation rather than recruitment of TNK2 is key to separate DEG- and REC-ligand signals, and could potentially be a key molecular switch in functional selectivity. TNK2 is a known clathrin-binding protein and has previously been reported to be involved with receptor traffic of EGFR and there are two inconsistent reports whether the knockdown of TNK2 leads to the reduced degradation of EGFR induced by EGF or not^38, 39^. Interestingly, Tahir et al. recently revealed an interaction between TNK2 and CLINT1^40^. Collectively, our integrated multilayered proteomics approach revealed a differentially regulated protein network consisting of CLINT1 SYNRG, and TNK2, which we decided to examine in more detail for their role in EGFR functional selectivity.

### CLINT1 regulates EGFR degradation

The CCV-related protein, CLINT1, also known as EPN4, is a member of the Epsin family in addition to EPN1, EPN2, and EPN3 with known functions in clathrin-mediated endocytosis (CME) including EGFR trafficking^33^. CLINT1 is structurally distinct within the family given the absence of binding domains for ubiquitin, AP2, and EPS15. Instead, CLINT1 has an AP/GGA motif and a methionine-rich domain. CLINT1 is also distinct in terms of function, and it is considered to be involved in transport from the trans-Golgi network to endosomes^34^. Sigismund et al. showed that knockdown of EPN1 and functionally related EPS15 and EPS15L1 reduced CME for EGFR. In our EGFR-TurboID proximity data, we quantified interaction profiles for EPN2, EPN3, CLINT1, EPS15 and EPS15L1, with CLINT1 showing the most pronounced difference between DEG- and REC-ligands (Fig. 5A). Although CLINT1 showed the highest abundance in the proteome data of the T1 HeLa clone, we deemed the proximal interaction with EGFR highly specific as the second highest ranking protein EPS15L1 did not show ligand-specific proximal recruitment profiles (Fig. 5A and Fig. 5B). This suggests that CLINT1 plays a critical role for the EGFR fate in response to REC-ligands.

**Figure 5.**
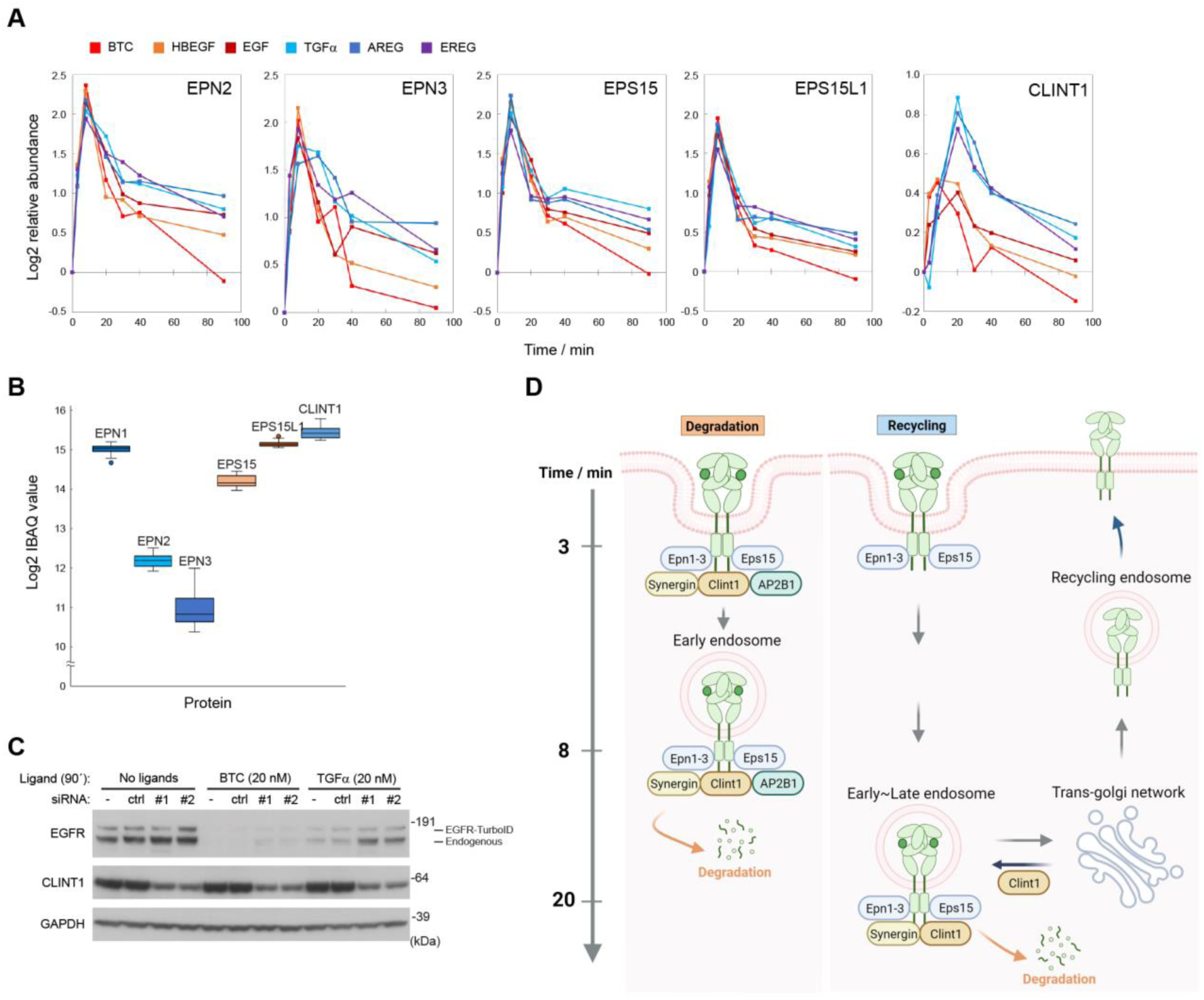
(A) Temporal profiles of log2 abundance of protein recruitments. The values are standardized by the value at 0 min. (B) Box plot showing log2 iBAQ values of Epsin family proteins, ESP15, and EPS15L1 (n=5). (C) Western blotting of the cell lysates of clone T1 cells stimulated with BTC (20 nM) or TGFα (20 nM) for 90 min after 48 h treatment with siRNA targeting CLINT1. (D) Hypothesis of functions of CLINT1 on receptor traffic of EGFR.

To investigate the involvement of CLINT1 and TNK2 in EGFR receptor fate, we evaluated ligand-induced EGFR degradation in CLINT1 or TNK2 knockdown cells. We found that EGFR degradation induced by BTC and TGFα was impaired by CLINT1 knockdown (Fig. 5C). In particular, the degradation rate was reduced under TGFα-stimulated conditions in the knockdown condition, indicating that CLINT1 contributes significantly to receptor degradation induced by recycling-inducing ligands. Considering the difference in the temporal recruitment profile of CLINT1, we hypothesized that the function of CLINT1 is different between ligand types. Upon activation by DEG-ligands, CLINT1 and AP2 are considered to interact with EGFR in AP2-positive clathrin-coated vesicles right after receptor activation. Our finding that CLINT1 knockdown impaired receptor degradation suggests that these CLINT1- and AP2-positive vesicles sort EGFR to lysosomes for degradation (Fig. 5D). Contrariwise, upon activation by REC-ligands, there is no AP2 recruitment but strong CLINT1 recruitment at later time points around 20-40 min. In this scenario, considering that trans-Golgi network is a pathway sorting receptors to recycling endosomes, CLINT1 could function to inhibit sorting of EGFR-positive late endosomes to trans-Golgi network, thereby causing degradation of the receptor (Fig. 5D).

### Behavior of CCV-related proteins induced by low EGF concentration and engineered chimeras of natural ligands

As our multilayered proteomics approach provided valuable insights into functional selectivity by quantifying intracellular signaling networks activated by EGFR ligands, we next focused on extracellular aspects of EGFR signaling by analyzing the effects of different ligand concentrations and engineered chimeras of natural ligands. Physiological concentrations of EGF vary widely ranging from picomolar to nanomolar, and ligand concentration can significantly influence EGFR signaling^15, 16^. Picomolar EGF mainly induces CME sorting receptors to the recycling pathway, while nanomolar EGF exceeds the capacity of CME to mainly induce clathrin-independent endocytosis (CIE) designating receptors to the degradation pathway. Thus, the ability to control ligand concentrations serves as a way to control EGFR fate. However, despite the wide use of recombinant growth factors in medical applications, the physiologic in vivo concentrations are difficult to control. Therefore, a more tractable alternative is engineering ligands with controlled signaling outputs at saturating concentrations.

Here, we designed four engineered ligand chimeras by combining the amino acid sequence of EGF and AR based on structural information for a region shared among all six natural ligands. This region comprises the consensus EGFR-binding motif (EGF-like domain) consisting of a beta-hairpin loop and a C-terminal region interacting with EGFR domain I and III, respectively (Fig. S5A) ^41^. For the chimera design, we were particularly interested in determining if this region is important for EGFR degradation and recycling. Thus, we divided the domain I-binding region into three parts (Figure S5A: 19–21, 25–30, and 32–38), and generated four chimeras (EGF/AR^19–21^, EGF/AR^25–30^, EGF/AR^32–38^, and EGF/AR^19–38^) by swapping the sequences of EGF to that of AR. HeLa cells were stimulated with various concentrations of the purified chimera ligands using wildtype EGF as positive control, and evaluated phosphorylation of EGFR and downstream signaling proteins by Western blotting (Fig. S5B). Based on this, we determined the saturating concentration for each chimera ligand (EGF/AR^19–21^: 20 nM, EGF/AR^25–30^: 500 nM, EGF/AR^32–38^: 50 nM, EGF/AR^19–38^: 100 nM), which were used in subsequent experiments. Next, we monitored the effect on EGFR levels by western blotting using lysates from cells stimulated with the four chimeric ligands for 90 min. EGF/AR^19–21^ showed EGFR levels similar to DEG-ligands, and noteworthy, EGF/AR^25–30^, EGF/AR^32–38^ and EGF/AR^19–38^ shifted the ‘original’ EGF response to mimic REC-ligands (Fig. 6A). As a result, we confirmed the ability of engineered chimeric ligands to shift EGFR fate and pinpointed a key minimal region in REC-ligands determining this feature.

**Figure 6.**
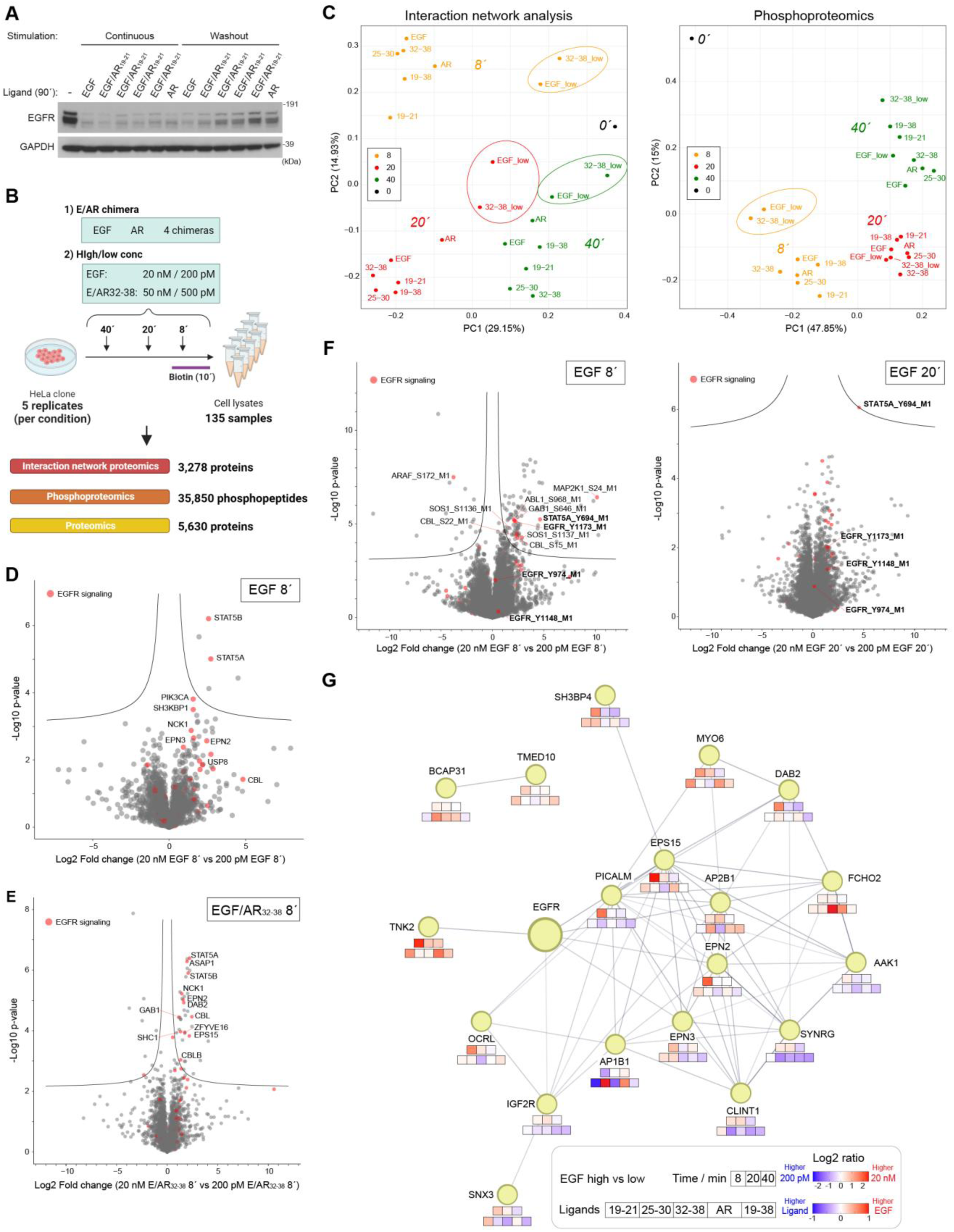
(A) Western blotting of the cell lysates of clone T1 cells stimulated with indicated ligands for 90 min. (B) Workflow of the multilayered proteomics analysis. (C) PCA depicting interaction network and phosphoproteomics data. (D) Volcano plots highlighting EGFR signaling-related proteins (red, KEGG term: “ErbB Signaling Pathway” and/or “Endocytosis”) from interaction network analysis data. Fold change represents high EGF (20 nM) treatment versus low EGF (200 pM) treatment for 8 min. (E) Volcano plots highlighting EGFR signaling-related proteins (red, KEGG term: “ErbB Signaling Pathway” and/or “Endocytosis”) from interaction network analysis data. Fold change represents EGF/AR^32-38^ (50 nM, 8’) treatment versus control. (F) Volcano plots highlighting phosphopeptides from EGFR signaling-related proteins (red, KEGG term: “ErbB Signaling Pathway”). Fold change represents high EGF (20 nM) treatment versus low EGF (200 pM) treatment for 8 or 20 min. (G) Interaction network of CCV-related proteins with their profiles of recruitment. The network is created using the STRING database.

Next, we performed an in-depth characterization of the intracellular effects of these chimeric ligands at multiple proteomics layers (interactome, phosphoproteome and proteome). We prepared lysates from HeLa clone T1 cells after stimulation with EGF/AR chimeras and for reference EGF and AR for 8, 20, or 40 min, and samples were analyzed according to our multilayered proteomics workflow (Fig. 6B). Moreover, to evaluate the effect of low and high ligand concentration on EGFR signaling, we included two different concentrations for EGF (200 pM and 20 nM) and EGF/AR^32–38^ (500 pM and 50 nM). We identified 3,278 proteins (no filtering) in the interaction network analysis, 35,850 phosphosites (Class I) in the phosphoproteome, and 5,630 proteins in the proteome analysis (Fig. 6B). The PCA plots showed separation by time and differences between low and high concentrations (Fig. 6C), and these effects were particularly pronounced in the interaction network data compared to phosphoproteome data showing only differences by concentration at 8 min stimulation. For the interaction network analysis, volcano plots of log2-fold changes (EGF: 20 nM versus 200 pM) and significance (–log10 (P-value)) showed a proximal preference for EGFR signaling-related proteins such as E3 ubiquitin ligase CBL and Epsins including a significant recruitment for STAT5A/B for the high concentration upon 8 min stimulation (Fig. 6D). These findings were also confirmed for EGF/AR^32–38^ (Fig. 6E). Upon 20 min stimulation with EGF, ESCRT-related proteins such as STAM, USP8, and VPS37 were preferentially recruited by high saturating concentration (Fig. S5C).

For the phosphoproteome and consistent with the PCA results, several phosphopeptides related to EGFR signaling were significantly and differentially regulated at 8 min for EGF high versus low concentration compared to 20 min stimulation (Fig. 6F) whereas EGF/AR^32–38^, showed significant phospho-regulation at both time points (8 and 20 min) (Fig. S5D). Noteworthy, only STAT5A phosphorylated at Y694 showed significantly increased levels for both ligands and time points (Fig. 6F and Fig. S5D) confirming regulation beyond recruitment to EGFR (Fig. 6F and Fig S5D). Interestingly, EGFR phosphorylated at Y1173 shared the same pattern of regulation. In contrast, we specifically detected for EGF/AR^32–38^ a difference in the number and level of concomitantly phosphorylated CBL phosphoserine peptides (Fig. S5D).

Finally, we extracted information on CCV-related proteins analogous to the analyses of the proximal proteins. In this dataset, 19 CCV-related proteins were quantified, and we mapped the log2 ratio of recruitment between high and low concentration of EGF (Fig. 6G, first row), and difference between each ligand chimera and EGF at 20 min (Fig. 6G, second row). Most proteins are recruited more strongly at high concentrations of EGF at 8 min and in favor of low concentrations of EGF after 20 min. Ligand chimeras that induce recycling were also found to strongly recruit proteins such as CLINT1, SYNRG, EPN2, and EPN3, which were implicated in ligand-dependent endocytic receptor fate.

### Tyrosine-mutated EGFR-TurboID reveals a site-specific comprehensive interaction network map and confirms STAT5 recruitment to EGFR pY1173

So far, we have shown that the intracellular interaction networks are differentially regulated depending on extracellular stimuli activating EGFR. Direct interactors of active EGFR associate with phosphorylated tyrosine sites on the EGFR C-terminal tail, which harbors nine tyrosine residues (number refers to immature/mature receptor: Y974/*Y998*, Y992/*Y1016*, Y1045/*Y1069*, Y1068/*Y1092*, Y1086/*Y1110*, Y1101/*Y1125*, Y1114/*Y1138*, Y1148/*Y1172*, and Y1173/*Y1197*). These nine sites differ in surrounding amino acid sequence inferring a motif-specific selectivity of recruited proteins (Fig. 7A). Information about site-specific recruitment of proteins and proximal complexes to individual tyrosine phosphorylation sites requires single phosphotyrosine site level resolution for EGFR to pinpoint where and how the interaction network changes.

**Figure 7.**
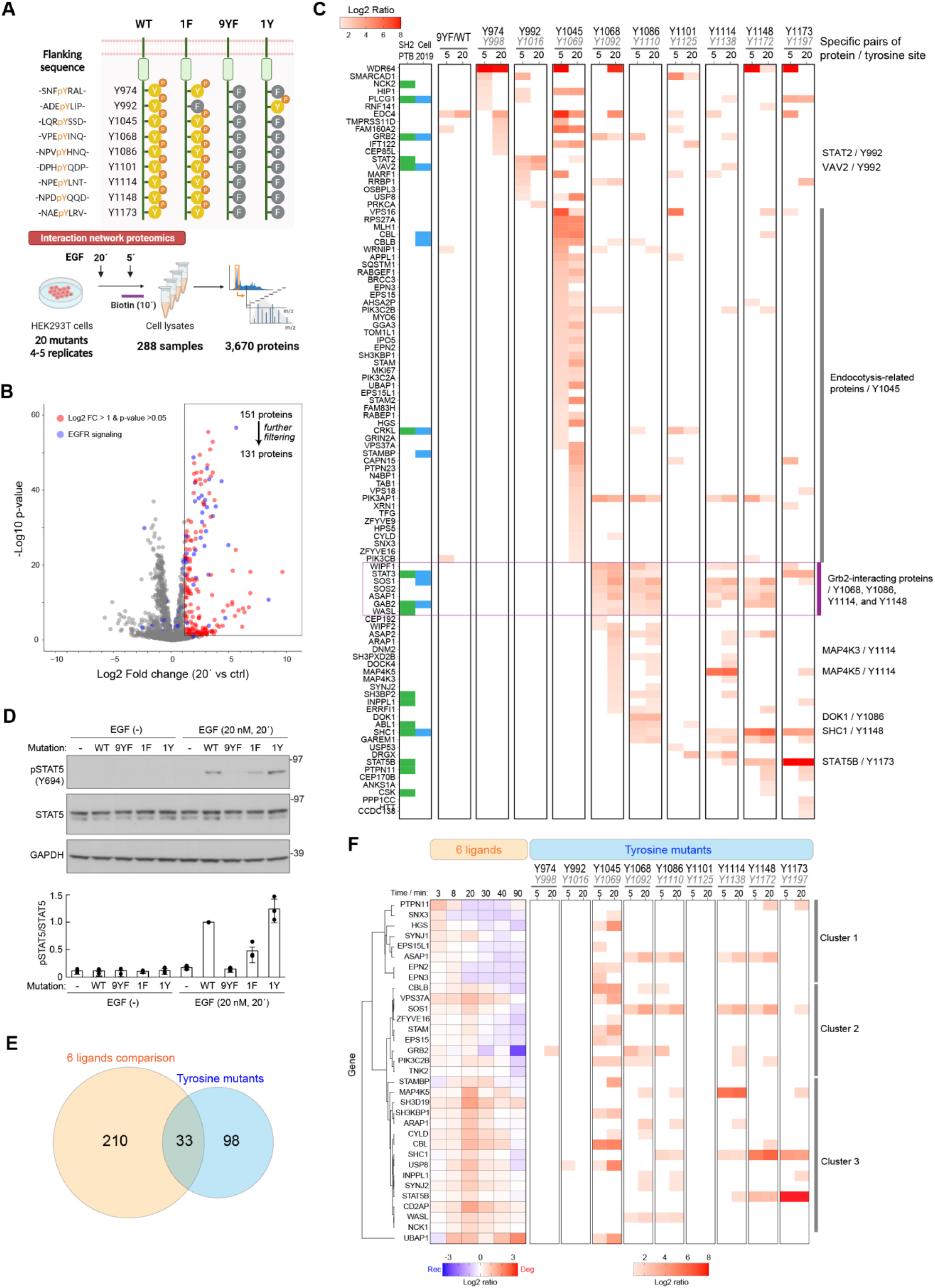
(A) Illustration of EGFR C-terminus tyrosine site, our design of tyrosine/phenylalanine mutants, and schematic illustration of the interaction network analysis. (B) Volcano plot highlighting regulated proteins between 20 min stimulation condition and control. (C) Heatmap showing the recruitment of the proteins to each tyrosine site. The ratio is calculated by subtracting the log2 value of 1F chimera from that of 1Y8F chimera and standardized by row-wise median subtraction. (D) Western blotting of the cell lysates of HEK293T cells expressing the WT, 9YF, Y1173F, or F1173Y mutant stimulated with EGF (20 nM) for 20 min. (E) Venn diagram showing the overlap of the identified proteins between 6 ligands data and tyrosine mutant data. (F) Heatmap integrating the information of the temporal recruitment profile and selectivity for tyrosine sites of 33 shared proteins.

To perform site-specific interaction network mapping in cells, we used our proximity labeling setup and designed a pair of EGFR-TurboID mutants for each tyrosine site (Fig. 7A): one with the tyrosine site mutated to phenylalanine (1F8Y), and one with the remaining eight tyrosine sites mutated to phenylalanine (1Y8F) to ultimately have 18 different mutants. Moreover, we designed an EGFR-TurboID mutant with all nine tyrosine sites mutated to phenylalanine (9YF) to allow for identification of potential background binders and recruitment to parts of EGFR independent of phosphorylated tyrosines. Comparison of these mutants (total 19) with wild type (wt) EGFR-TurboID would allow for the site-specific mapping of interaction networks. Given the comprehensiveness of this panel, we used a transient expression system in HEK293T cells. We initially transfected these cells with plasmid DNA encoding wtEGFR-TurboID and confirmed a 3-fold higher expression level compared to endogenous EGFR by western blotting (Fig.S6A). Moreover, EGF stimulation resulted in EGFR and EGFR-TurboID phosphorylation and downstream ERK1/2 activation. Flow cytometry analysis confirmed an approximately 50% transfection efficiency and 20 min stimulation with EGF reduced the surface expression level of EGFR-TurboID due to the internalization (Fig. S6B). From these results, we concluded that the molecular behavior of the transiently modestly overexpressed EGFR-TurboID aligned well with that of endogenous EGFR and hence, we deemed this setup suitable for our comprehensive site-specific mapping approach.

First, we evaluated impact on protein recruitment caused by the overexpression of EGFR-TurboID, and performed the interaction network proteomics analysis using lysates from non-transfected HEK293T cells and wtEGFR-TurboID-expressing cells with or without EGF stimulation. Comparing non-transfected cells and transfected cells, Several EGFR signaling-related proteins are already overrepresented in the transfected cells even without EGF treatment (Fig. S6C). This result implies that overexpressed EGFR-TurboID causes EGF-independent interaction with several proteins. However, by comparing the transfected cells with and without EGF treatment, we can clearly confirm the enrichment of the EGFR signaling-related proteins including ones showing background recruitment (Fig. S6D).

For large-scale proximity interaction network proteomics, we stimulated cells expressing one of the EGFR-TurboID mutants with 20 nM of EGF for 5 or 20 min, and used the resulting cell lysates for SA/biotin enrichment and LC-MS/MS analysis (Fig. 7A). We identified and quantified 3,670 proteins from all the pull-down samples (Table S7). We focused our subsequent analysis on proteins identified to associate with EGFR-TurboID under minimally perturbed conditions including wtEGFR-TurboID and 1F8Y mutant data upon EGF stimulation. From a volcano plot analysis of log2-fold change of 20 min EGF stimulation versus control and significance (–log10 (P-value), we selected 151 proteins showing >two-fold change upregulation and P-value less than 0.05 (shown in red in Fig.7B, Fig. S6E). We further narrowed down the number of proteins by other filtering steps (see Methods section). From a list of 131 candidate proximal proteins represented in the heatmap in Fig. S6E, we interpreted binding of a protein to a specific site upon intensity increase for 1Y8F mutant data. Additionally, if the binding is phosphotyrosine-specific, the intensity from corresponding 1F8Y mutant data should decrease as a consequence of abolished phospho-tyrosine interaction. We summarized specific recruitment to a certain site by visualizing the Log2-fold change of the intensity of paired 1Y8F mutant versus 1F8Y mutant in a heatmap for 5 and 20 min EGF stimulation compared to control (Fig. 7C). Based on this, we identified highly selective protein-phosphotyrosine site pairs, such as STAT2 (Y992), DOK1 (Y1086), MAP4K5 (Y1114), CBL (Y1045), SHC1 (Y1148/Y1173), PLCγ (Y1173), and STAT5 (Y1173). While the EGFR specific binding sites of DOK1^42^, CBL^43^, SHC1^44^, and PLCγ^45^ are consistent with previous reports, those of STAT2, MAP4K5, and STAT5 have not previously been confirmed. The canonical CBL-binding Y1045 site is critical for receptor internalization^43^, and the data indeed showed many proteins related to this process binding to this site with high specificity (highlighted in purple in Fig. S7). It is well known that once the receptor is activated, CBL binds to pY1045 and ubiquitinates the EGFR C-terminal tail triggering the assembly of endosomal proteins. In agreement with previous reports, we confirmed CBL binding to pY1045 with high specificity in addition to a range of proteins with similar profiles including ubiquitin protein RPS27A, ESCRT-related proteins (USP8, STAM, VPS37A, UBAP1), endocytosis-related proteins (HIP1, HIP1R, MYO6, ARPC1B, and EPS15) and the deubiquitinase BRCC3.

Proteins can show some degree of binding redundancy to different phosphotyrosine sites if they bind directly or as part of larger protein complexes. However, given our approach based on proximity labeling, we speculated that a lack of increase in 1Y8F/1F8Y ratios could be a consequence of biotinylation of the protein when bound to neighboring sites alone or as part of a complex for the 1F/8Y mutant. In this case, we would not detect a ratio increase and deemed the proximal protein specific to the phosphotyrosine site. Thus, to improve our site-specific protein map, we corrected for this potential redundancy and normalized the ratio by subtracting median for each protein (Fig. 7C). After this data normalization, four tyrosine sites (Y1068, Y1086, Y1114, and Y1148) showed similar overall protein recruitment (highlighted in purple in Fig. 7C). Of note, these four sites are known Grb2 binding sites^46, 47^, which we confirmed for 3 of the four sites. Grb2 engages with EGFR in larger protein complexes with proteins related to Ras-MAPK pathway regulation including SOS1, SOS2, and Gab2, which we also confirmed here for all four sites.

Among the protein-tyrosine site pairs discovered, the EGFR pY1173-STAT5 interaction is of particular interest because the specific EGFR recruitment site for STAT5 has been a controversial topic. Kloth et al. proposed Y845 as STAT5 binding site as a Y845F mutation largely downregulated the activation of STAT5^48^. However, Schulze et al. showed that STAT5 binds to phosphopeptides derived from Y954 region or Y974 region, but not from Y845 region^47^. Here, we show STAT5 recruitment to site Y1173 and confirmed by western blotting activating Y694 phosphorylation of STAT5 by EGF (Fig. 7D). While the 9YF mutant did not induce the Y694 phosphorylation of STAT5, the F1173Y mutant activated STAT5 even stronger than WT EGFR. This result implies that pY1173 site functions as a direct recruitment site for STAT5 and is critical for STAT5 activation. Furthermore, this is in agreement with the observation that high concentrations of EGF induced a distinct phosphorylation of the EGFR Y1173 site and STAT5 Y694 (Fig. 6E).

Finally, we compared the proximal interaction network data from the six-ligand and tyrosine mutant analyses to determine the contribution of each tyrosine residue to functional selectivity. The six-ligand comparison revealed 243 proteins that differed substantially between the two ligand groups and 131 proteins were the focus of the tyrosine mutant analysis. We found that 33 proximal proteins were common among the datasets (Fig. 7E), and performed clustering of them based on the difference of the recruitment profiles between the two ligand types, and integrated the information of their selectivity to different EGFR tyrosine residues (Fig. 7F). Most of these 33 proteins were more strongly regulated by the DEG-ligands immediately after stimulation, and then either more strongly regulated by the REC-ligands after about 20 minutes (cluster 1), more strongly regulated by the REC-ligands after 40 to 90 minutes (cluster 2), or more regulated by DEG-ligands over time (cluster 3). Many of the proteins in clusters 1 and 2 are recruited to Y1045, consistent with this site being the most important residue controlling endocytosis linked to receptor fate. On the other hand, proteins such as CBL, USP8, and SH3BP1 are Y1045-dependent but more strongly oriented toward DEG-ligands, suggesting that subtle differences in the distribution of protein abundances in the Y1045-centered interactome determine receptor fates.

## Discussion

This study provides integrated data of interaction networks and phosphoproteomes serving as a huge resource of the molecular network dynamics of EGFR signaling. Despite several previous studies on the analysis of the EGFR interactome based on the proximity-dependent labeling technique^20-22^, our study offers several advantages over these. Firstly, we showed the importance of having fine-tuned EGFR expression to physiological level to study endocytic receptor trafficking. Although overexpression of EGFR-fusion constructs improves the sensitivity of the MS-based detection, the natural endocytic behavior of EGFR is distorted, which makes it difficult to draw conclusions on differential behavior of different ligands and functional selectivity. Secondly, EGFR signaling is governed not only by protein-protein interactions but also by phosphorylation cascades, so it is necessary to combine more than one layer of information to elucidate complex mechanisms of cell signaling. Here, we combined proximity labeling-based interaction networks with phosphoproteomics to study EGFR signaling and trafficking with unprecedented spatiotemporal resolution. Lastly, we have extended the knowledgebase on EGFR functional selectivity by profiling all six highest-affinity ligands offering comprehensive comparative layers of signal information. Previous studies have been limited to the analysis of only one or two ligands, usually EGF alone or in comparison to TGFα. However, to meet the demand of high-throughput studies, we developed a semi-automated, high-throughput MS-based proteomics approach. This enabled us to analyze the six ligands with the temporal resolution required to track receptor trafficking, including a sufficient number of biological replicates needed for robust MS-based label-free quantification, ultimately comprising >1,500 samples for the comprehensive study of EGFR signaling diversity.

By the comparative study of ligand grouped into degradation- and recycling-inducing ligands, we found that clathrin coated vesicle-related proteins including clathrin interactor CLINT1, SYNRG, AP2B1 showed different temporal profiles of their recruitment to EGFR. These findings are specifically supported by our time-resolved and quantitative proteomics approach. We demonstrated that the balance of EGFR endocytic fate can be biased to receptor recycling by reducing the expression level of CLINT1, revealing its role as a molecular EGFR trafficking switch.

The study of interaction partners to activated EGFR involves the analysis of proteins binding to phosphorylated tyrosine sites in the C-terminal tail. To date, tyrosine mutation analysis of EGFR to pinpoint phosphorylation site-dependent protein interactors has been performed in two ways: either by co-immunoprecipitation of known adapter proteins in EGFR wild type or under tyrosine-mutated conditions^42-46^ or by MS-based analysis of pull-downs using phosphotyrosine-containing peptide baits^47^. Phosphopeptide-based pull-downs are limited to the detection of proteins interacting with a single phosphotyroinse site, and can be challenged by non-specific bindings as bait peptides exist outside their natural conformational environment. Moreover, these analyses lose spatial information due to the pull down procedure in cell lysates. Here, we applied our high-throughput interaction network analysis based on proximity labeling to systematically evaluate the selectivity of proteins binding to individual phosphorylated tyrosine residues in the C-terminal tail of EGFR. This method offers the advantage of studying full-length EGFR expressed in cells with labeling of proximal proteins in intact cells. We mapped the selectivity of all nine tyrosine residues in the C-terminal tail using a complete panel of phosphosite-dead tyrosine-to-phenylalanine mutated EGFR-TurboID constructs. Using a smart design of preparing a pair of EGFR mutants for each site, one with the tyrosine site mutated to phenylalanine (1F8Y), and one with the remaining eight tyrosine sites mutated to phenylalanine (1Y8F). We performed pairwise comparison of protein recruitment to discover site-specific interactors in a ligand-dependent manner. Reassuringly, we identified a large part of the known phosphotyrosine site-binding interactors. Moreover, we revealed the specific binding of STAT5 to EGFR pY1173, which was not known before. This validates our finding that EGFR pY1173 and STAT5 pY694 are primary co-regulated sites when comparing high (degradation-inducing) versus low (recycling-inducing) concentrations of EGF stimulation. Collectively, our comprehensive dataset provides a vast resource of molecular networks of EGFR signaling for a deeper understanding of the molecular mechanisms underlying functional selectivity.

## Figures

**Figure S1.**
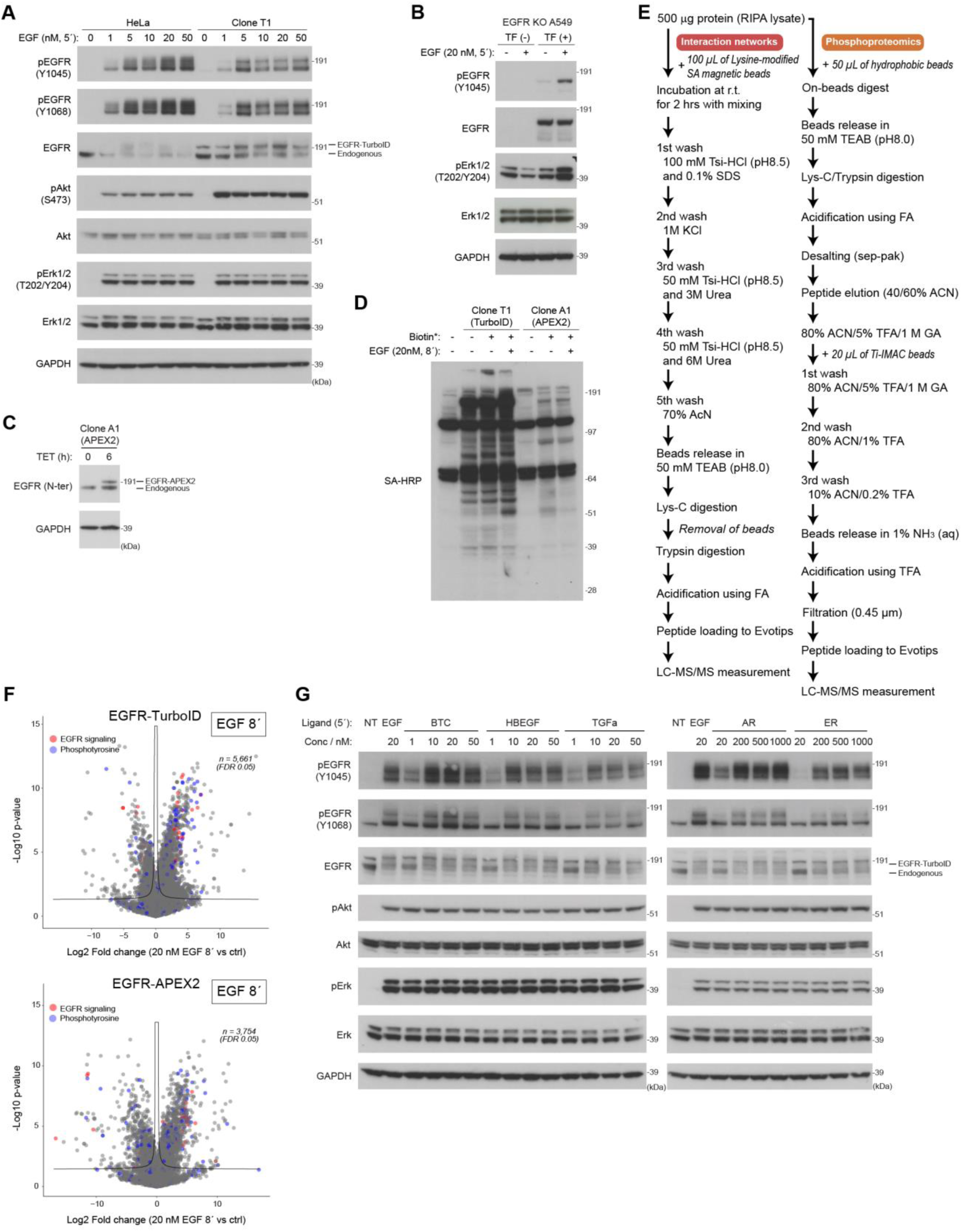
(A) Western blotting of the cell lysates of HeLa and clone T1 cells stimulated with EGF (20 nM) for 5 min. (B) Western blotting of the cell lysates of EGFR-TurboID-expressing A549 cells (EGFR-knockout) stimulated with EGF (20 nM) for 5 min. (C) Western blotting of the cell lysates of APEX2 clone A1 treated with TET for 6 h. (D) Western blotting of the cell lysates of Clone T1 and APEX2 clone A1 stimulated with EGF (20 nM) for 5 min. APEX2 clone A1 cells were treated with TET for 6 h prior to the ligand stimulation. (E) Brief protocol of sample preparation for interaction network and phosphoproteomics analysis (F) Volcano plots highlighting peptides in EGFR signaling-related proteins (red, KEGG term: “ErbB Signaling Pathway”) and tyrosine-phosphorylated peptides (blue) from phosphoproteomics data using TurboID. Fold change represents EGF (20 nM, 8’) treatment versus control. (G) Western blotting of the cell lysates of clone T1 cells stimulated with indicated ligands at indicated concentrations for 5 min.

**Figure S2.**
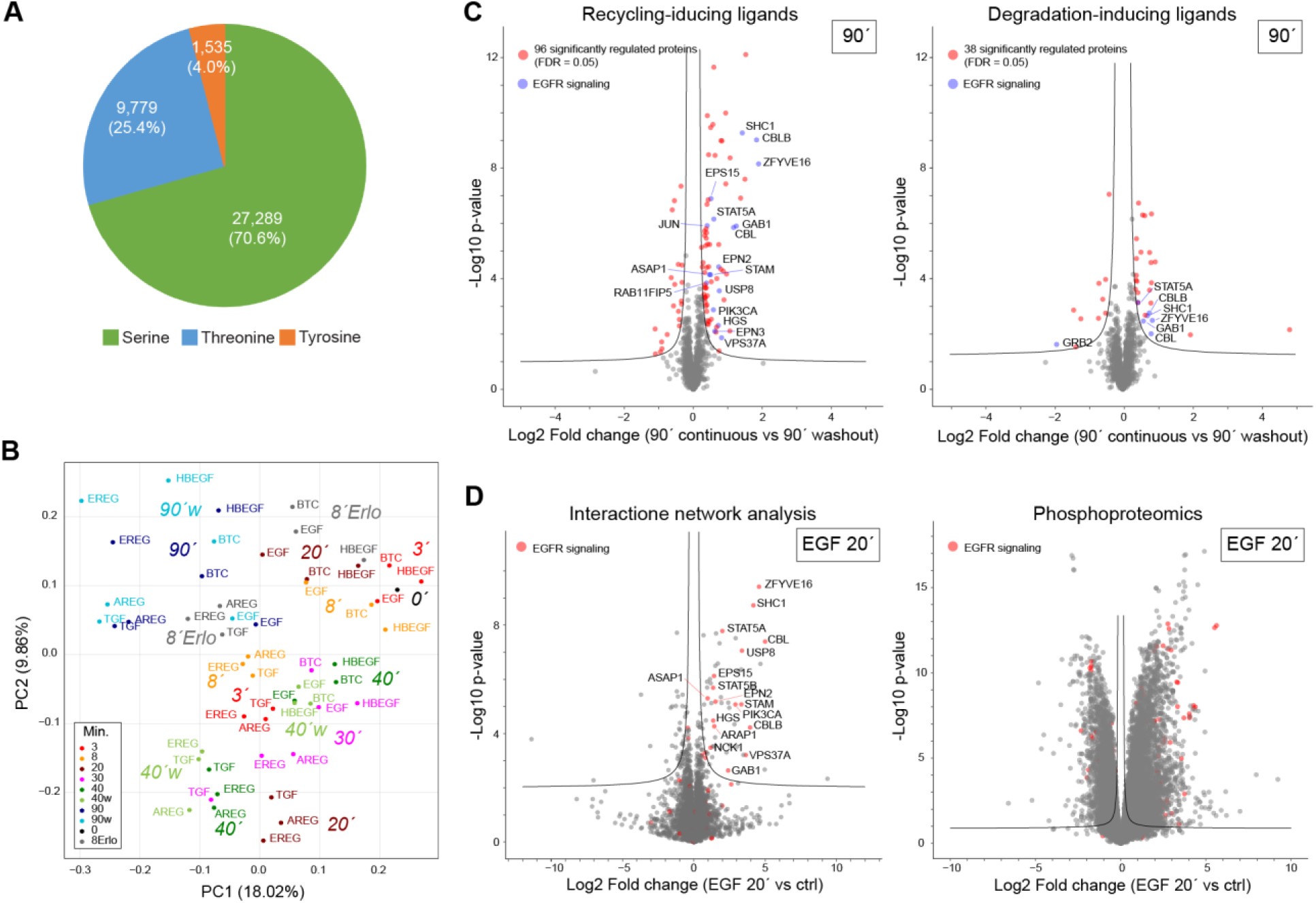
(A) Distribution of phosphorylated amino acids from phosphoproteomics data. (B) Volcano plots showing differentially regulated proteins from interaction network analysis data or phosphopeptides from phosphoproteomics data. Fold change represents 20 min stimulation with EGF versus control. (C) PCA of the log2-transformed abundance from interaction network analysis data. (D) Volcano plots showing differentially regulated proteins (red) for each ligand group, highlighting EGFR signaling-related proteins (blue, KEGG term: “ErbB Signaling Pathway” and/or “Endocytosis”). Fold change represents 90 min stimulation with washout method versus continuous method.

**Figure S3.**
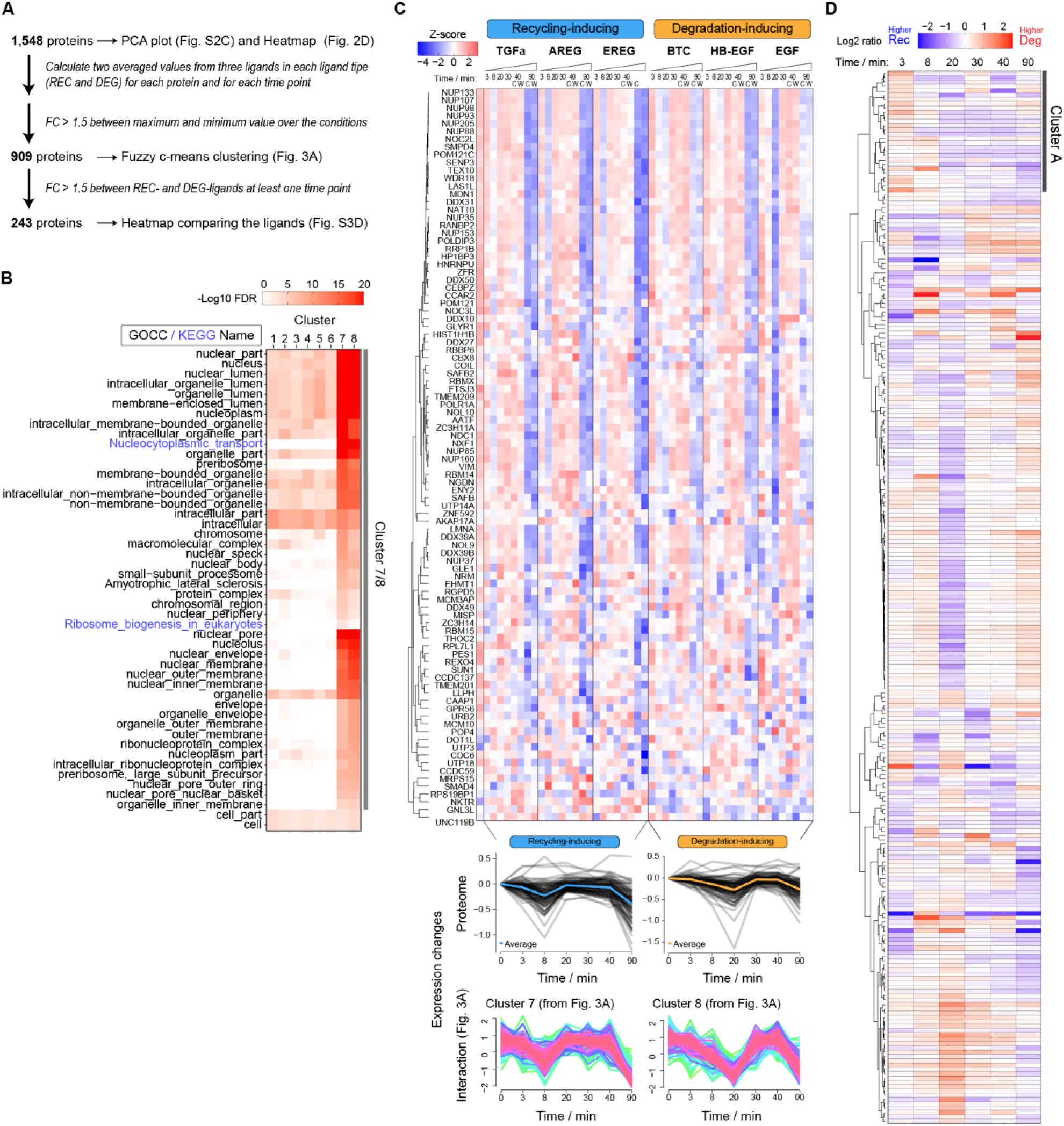
(A) Filtering steps of proteins identified in interaction network analysis data. (B) GOBP and KEGG enrichment analysis based on the proteins in each cluster. (C) Heatmap based on log2 abundance of proteins that are assigned in cluster 7 for recycling-inducing ligands and in cluster 8 for degradation-inducing ligands. (D) Heatmap of the proteins differently regulated by two types of ligands. The color scale represents the difference between recycling-(blue) and degradation-(red) inducing ligands. Median value from three ligands in each group was used for calculating the ratio.

**Figure S4.**
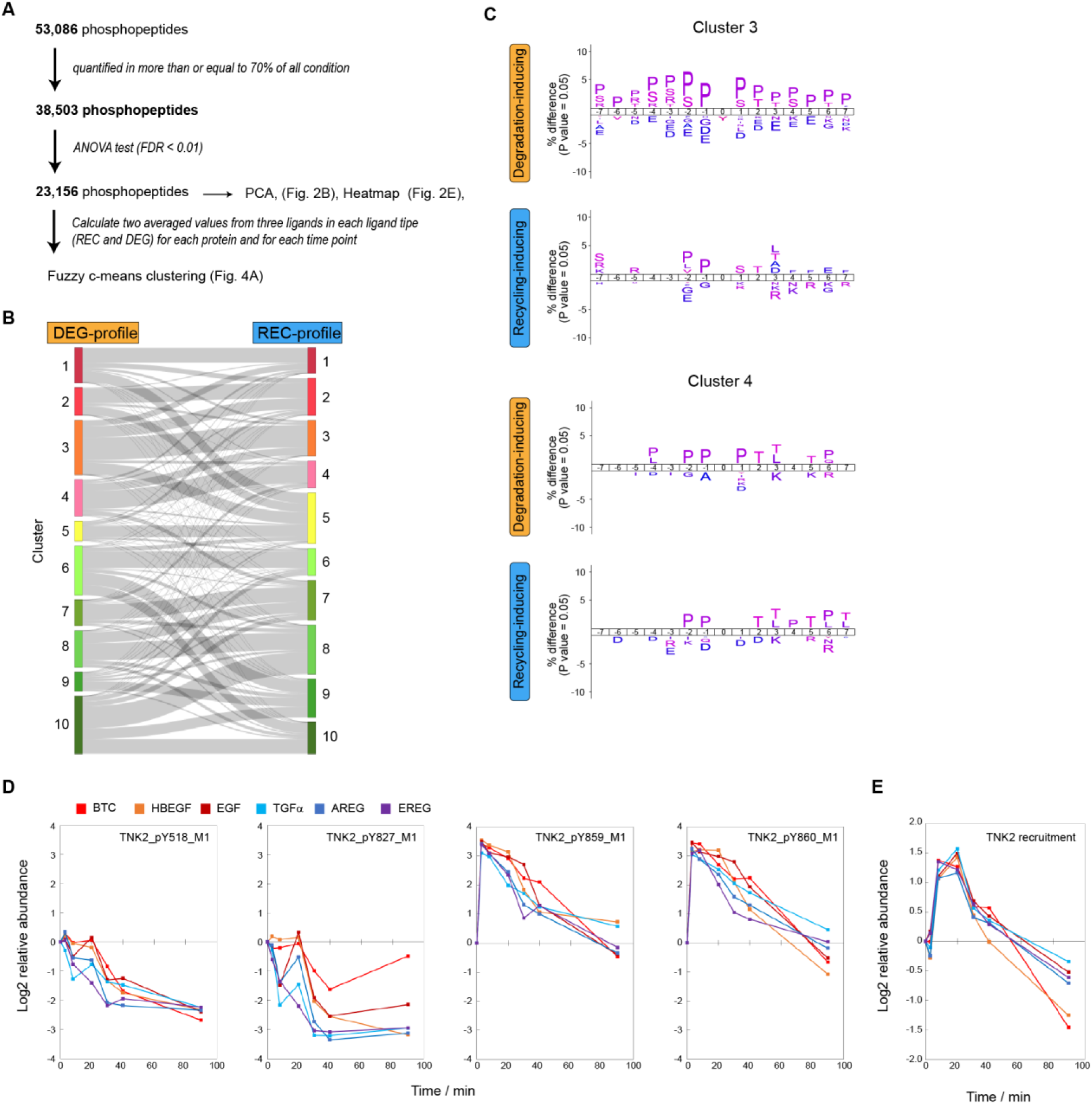
(A) Filtering steps of phosphopeptides identified in phosphoproteomics data. (B) Sankey diagram describing the association of clusters between degradation- and recycling-inducing ligands. (C) Enriched phosphosites analysis using 15 sites surrounding phosphosites in clusters 3 and 4. The figures are created by IceLogo. (D, E) Temporal profiles of log2 abundance of (D) TNK2-derived phosphopeptides or (E) TNK2 recruitment. The values are standardized by the value at 0 min.

**Figure S5.**
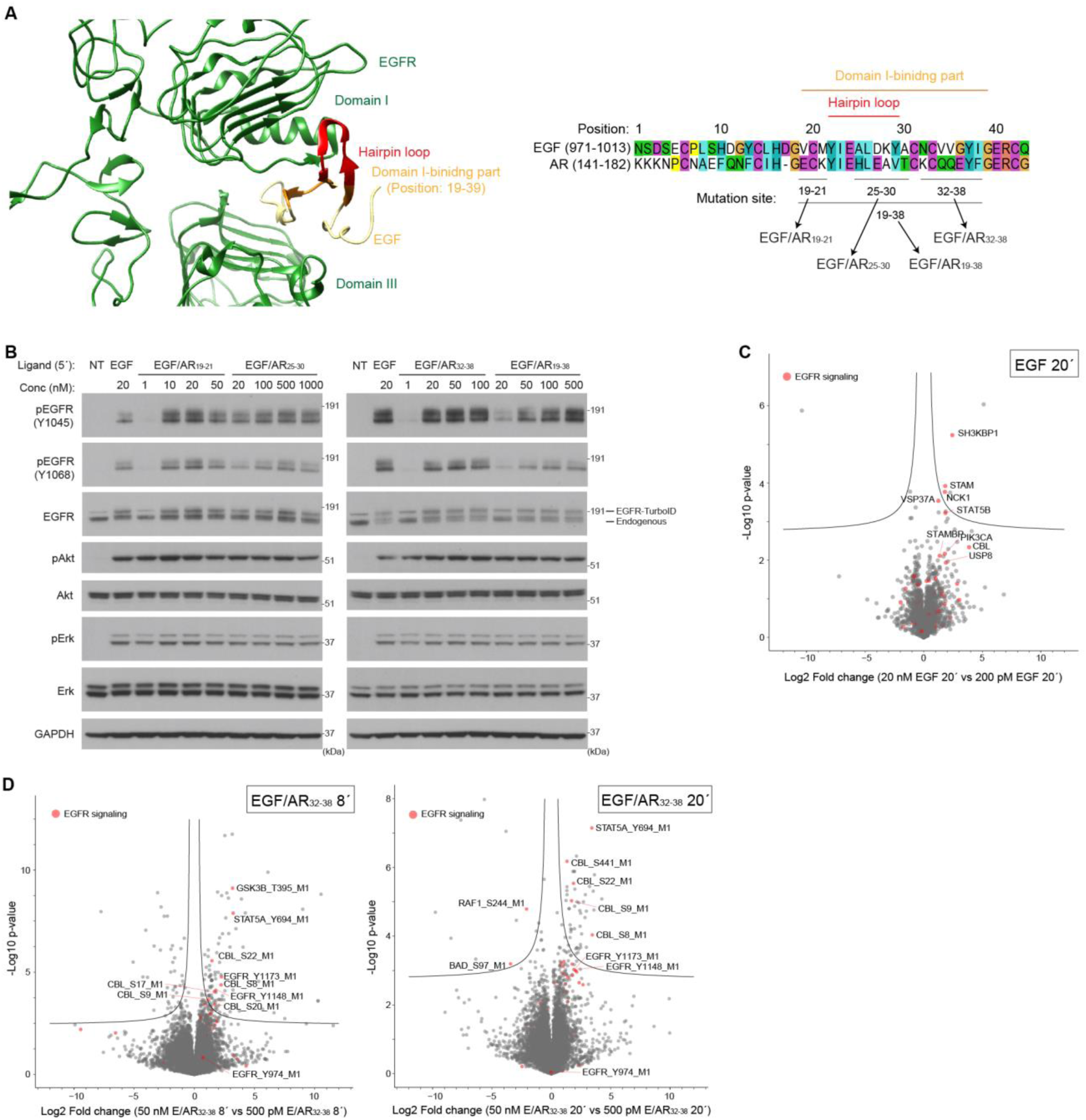
(A) Crystal structure of EGFR-EGF complex (Uniprot ID: 1IVO) and amino acid sequence of EGF-like domain of EGF (P01133: aa971–1013) and AR (P15514: aa141–182). (B) Western blotting of the cell lysates of clone T1 cells stimulated with indicated ligands at indicated concentrations for 5 min. (C) Volcano plots highlighting EGFR signaling-related proteins (red, KEGG term: “ErbB Signaling Pathway” and/or “Endocytosis”) from interaction network analysis data. Fold change represents EGF (20 nM, 20’ or 40’) treatment versus control. (D) Volcano plot highlighting phosphopeptides from EGFR signaling-related proteins (red, KEGG term: “ErbB Signaling Pathway”) from phosphoproteomics data. Fold change represents EGF/AR^32-38^ (50 nM, 8’ or 20’) treatment versus control.

**Figure S6.**
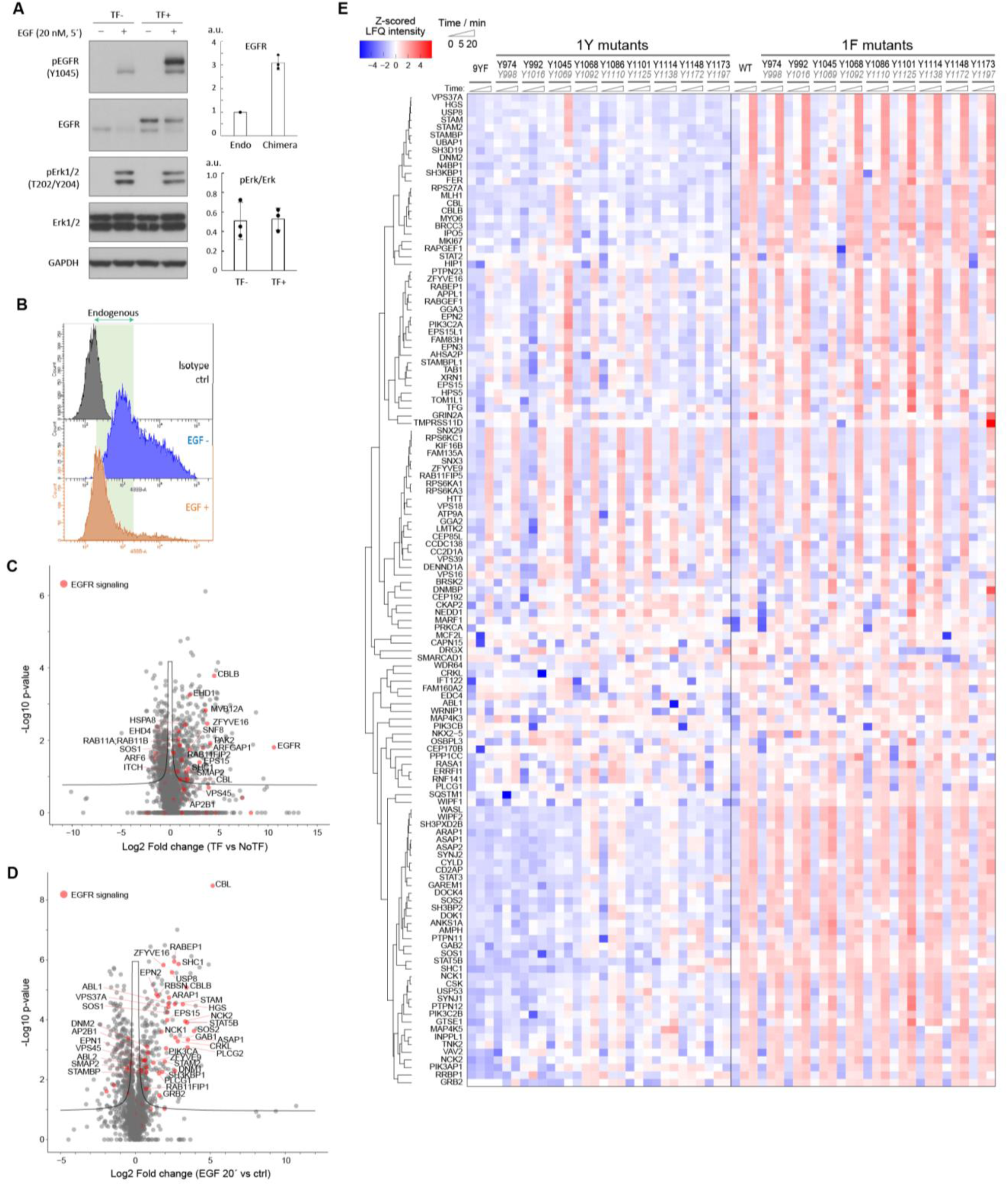
(A) Western blotting of the cell lysates of clone T1 cells stimulated with EGF (20 nM) for 5 min. The cells were transfected with EGFR-TurboID-expression vector 48 h before the stimulation. Values are mean ± s.d. from three independent experiments. (B) Flow cytometry analysis showing the surface expression level of EGFR after 20 min stimulation with EGF (20 nM). The cells were transfected with EGFR-TurboID-expression vector 48 h before the stimulation. (C, D) Volcano plots highlighting EGFR signaling-related proteins (red, KEGG term: “ErbB Signaling Pathway” and/or “Endocytosis”). (C) Fold change represents EGFR-Turbo-expressing condition and control. (D) Fold change represents 20 min stimulation with EGF (20 nM) versus control. (E) Heatmap of 131 proteins after the filtering steps.

**Figure S7.**
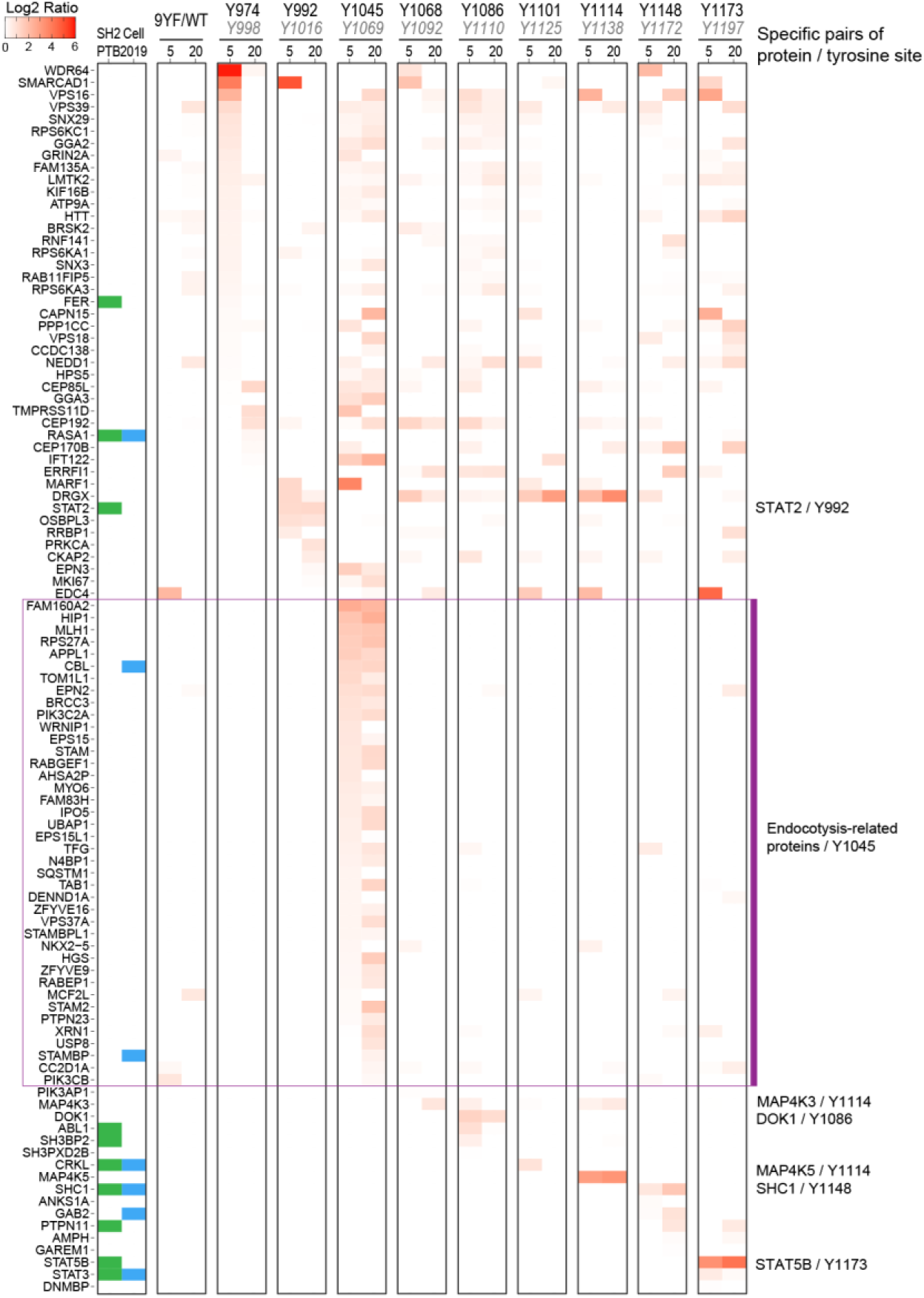
Heatmap showing the recruitment of the proteins to each tyrosine site. The ratios are calculated by subtracting the log2 values of 1F8Y chimera from those of 1Y8F chimera data.

## Material and methods

### Cell culture

HeLa, HeLa Flp-in T-REx, T1, A549 (EGFR KO)^1^, HEK293T cells were maintained at 37 °C in a humidified atmosphere with 5% CO2 in DMEM supplemented with 10% heat-inactivated fetal bovine serum (FBS) and penicillin/streptomycin (100 units/mL and 100 mg/mL; Thermo Fisher Scientific). The medium without FBS was used as starvation medium.

### Plasmids

We generated the construct: pcDNA3.1-EGFR-flag-APEX2 by amplifying the sequence encoding EGFR from the plasmid pEGFP-N1-EGFR-EGFP (Addgene: #32751) and cloning it into pcDNA3.1 (+) hygro BRCA1-FLAG-APEX2 (a kind gift from Rajat Gupta, Choudhary group, University of Copenhagen) substituting BRCA1. From pcDNA3.1-EGFR-flag-APEX2, we generated pcDNA3.1-EGFR-flag-TurboID by substituting APEX2 with a GeneStrands fragment (Eurofins) encoding TurboID (Addgene: #124646). From these vectors EGFR-flag-APEX2 and EGFR-flag-TurboID were subcloned into pcDNA5/FRT/TO to generate pcDNA5/FRT/TO-EGFR-flag-APEX2 and pcDNA5/FRT/TO-EGFR-flag-TurboID, which were used for Tet-inducible expression. For the phosphotyrosine mutation study, the enhancer sequence of pcDNA3.1 was shortened in order to reduce the expression level of EGFR-TurboID. For the 9YF mutant, the amino acid sequence corresponding to the C-terminal tail of EGFR was swapped to the sequence carrying nine phenylalanine mutations. The vectors for 1Y and 1F mutants were prepared by introducing point mutations to a wildtype vector and a 9YF vector, respectively.

For protein expression and purification of the ligand chimeras: The sequence of EGF was subcloned by LIC cloning into a pET expression vector with His6 and GST tag at the N-terminal fusion, followed by TEV protease cleavage site. Mutations were introduced by swapping indicated sequences of EGF to the corresponding sequence of AR.

### Protein purification

SHuffle competent cells were transformed with an expression vector. Cells were cultured at 30 °C, and protein expression was induced by supplementing Isopropyl β-D-1-thiogalactopyranoside (IPTG) (0.5 mM) when optical density reaches 0.6. After overnight incubation, cells were collected by centrifuge, sonicated in binding buffer (20 mM Tris-HCl (pH 8.0), 150 mM NaCl, 20 mM imidazole, 0.5 mM tris(2-carboxyethyl)phosphine (TCEP), cOmplete™ Protease Inhibitor Cocktail (Roche), and benzonase), and supernatants were collected by centrifuge. The supernatant was purified using a Ni-NTA Column (Cytiva). The eluted sample was supplemented with TEV protease and dialyzed against 20 mM Tris-HCl (pH 8.0), 500 mM NaCl, and 5 mM imidazole overnight. The dialyzed sample was applied to a Ni-NTA column, and the flowthrough was purified using a Superdex75 column (Cytiva). The concentration of the protein was determined by measuring the absorbance at 280 nm using a NanoDrop ND-2000 (Thermo Fisher Scientific). The purity of the final preparation was checked by sodium dodecyl sulfate-polyacrylamide gel electrophoresis (SDS-PAGE) and Coomassie Brilliant Blue staining.

### Cell lysis and western blotting

HeLa cells (2.0 × 10^5^ cells) or Clone T1 cells (4.2 × 10^5^ cells) were seeded in 6-well plates, and next day the medium was replaced by starvation medium and cultured 16–24 h. After the starvation, the cells were stimulated with the indicated concentration of ligand samples for indicated time at 37 °C. For the washout method, medium was removed 20 min after the stimulation with ligands, washed twice with pre-warmed starvation medium, and replenished with pre-warmed starvation medium. After the stimulation, the cells were washed with ice-cold PBS and lysed using modified RIPA buffer (50 mM Tris-HCl (pH7.5), 150 mM NaCl, 1% NP-40, 0.1% sodium deoxycholdate, 1 mM ethylenediaminetetraacetic acid (EDTA), 5 mM β-glycerophosphatase, 5 mM NaF, 1 mM sodium orthovanadate) supplemented with protease inhibitor cocktail. The buffer was supplemented with 0.1% SDS only when phosphorylation of STAT5 was analyzed. After the incubation on ice for more than 10 min, the lysates were scraped and centrifuged at 12,000 × *g* for 20 min at 4°C, and the supernatants were collected. The protein concentration of the cell lysates was measured using BCA assay kit and adjusted so that the amounts of proteins were the same between samples analyzed on the same gel. The cell lysates were mixed with NuPAGE LDS Sample Buffer (4×) (Thermo Fisher Scientific) supplemented with 100 mM DTT, boiled for 10 min at 70°C, separated by SDS-PAGE, and transferred to PVDF membranes. After the transfer, the membranes were blocked with 5% BSA/PBS-T, incubated with indicated primary antibody overnight at 4°C, and incubated with HRP-conjugated secondary antibody for 1h at ambient temperature, and developed using Novex™ ECL Chemiluminescent Substrate Reagent Kit (Thermo Fisher Scientific). Signal intensity was quantified using ImageJ. Primary antibodies used were rabbit anti-EGFR (ab32198; Abcam), rabbit anti-phospho-EGFR (pY1045) (#2237; Cell Signaling Technology), mouse anti-phospho-EGFR (pY1068) (#2236; Cell Signaling Technology), rabbit anti-BirA (NBP2-59939; Novus Biologicals), rabbit anti-Akt rabbit (#9272; Cell Signaling Technology), rabbit anti-phospho-Akt (S473) (#9271; Cell Signaling Technology), anti-Erk1/2 (#4695; Cell Signaling Technology) mouse anti-phospho-Erk1/2 (T202/T204) (#9106; Cell Signaling Technology), rabbit anti-Clint1 (ab251892; Abcam), rabbit anti-STAT5 (#9363; Cell Signaling Technology), rabbit anti-phospho-STAT5 (Y694) (#4322; Cell Signaling Technology), and mouse anti-GAPDH (Abcam).

### Cell lysis (APEX samples)

Clone A1 cells (3.0 × 10^5^ cells in 6-well plates or 2.7 × 10^6^ cells in P15 dishes) were seeded and cultured for 2 days. Then the medium was replaced by starvation medium supplemented with tetracycline (1 µg/mL). After 6h of starvation, the cells were incubated in starvation medium supplemented with Biotin-Phenol (500 µM) for 30 min, followed by stimulation with EGF (20 nM) for 8 min at 37 °C. H2O2 (1 mM) was added 1 min before harvest. After the stimulation, the cells were washed subsequently with quenching buffer (10 mM sodium azide, 1 M sodium ascorbate, and 0.5 M 6-hydroxy-2,5,7,8-tetramethylchroman-2-carboxylic acid in PBS) and ice-cold PBS, and lysed using modified RIPA buffer supplemented with protease inhibitor cocktail. After the incubation on ice for more than 10 min, the lysates were scraped and centrifuged at 12,000 × *g* for 20 min at 4°C, and the supernatants were collected. The protein concentration of the cell lysates was measured using Bradford Protein Assay kit and adjusted so that the amounts of proteins were the same between samples analyzed on the same gel.

### Transfection and RNA interference

EGFR-knockdown A549 cells (2.0 × 10^5^ cells), HEK293T cells (2.0 × 10^5^ cells), or T1 cells (3.0 × 10^5^ cells) were seeded in 6-well plates, and next day the cells were transfected with 1 µg of the expression vector using Lipofectamine 3000 reagent or 25 pmol of double-stranded siRNA using Lipofectamine RNAiMAX reagent. On the day after the transfection, the medium was replaced by starvation medium and cultured 16–24 h. Cell lysis and western blotting were performed as described in “Cell lysis and western blotting”. The double-stranded siRNA oligonucleotides targeting Clint1 were purchased from eurofin and their sequences are 5’-AAUACAGAUAUGGUCCAGAAA-3’ (called siRNA#1) and 5’-GCUCCUAGCUUACCUCAU A-3’ (called siRNA#2). Stealth RNAi™ siRNA Negative Control, Med GC (Thermo Fisher Scientific) was used as a negative control.

### Confocal microscopic imaging

T1 cells (2.0 × 10^4^ cells) were seeded in 8-well chamber slides, and next day the medium was replaced by starvation medium. After overnight starvation, cells were stimulated with EGF (20 nM) for 8 min at 37 °C. Biotin was added 10 min before the following fixation process. After the stimulation, cells were washed with PBS, fixed with 4% paraformaldehyde in PBS for 10 min at ambient temperature, and permeabilized with 0.2% TritonX-100/PBS for 5 min at ambient temperature. Then cells were washed twice with PBS, blocked with 1% BSA in PBS-T for 30 min at ambient temperature, washed twice with PBS, incubated with primary antibody in 1% BSA in PBS-T for 1h at ambient temperature, washed twice with PBS, incubated with secondary antibody in 1% BSA in PBS-T for 1h at ambient temperature. The nuclei part was stained with DAPI. All the images were acquired at room temperature with a confocal microscope LSM880 (Zeiss). Primary antibodies used were mouse anti-EGFR antibody (#GR01; Merck Millipore), goat anti-EEA1 (#sc-6414, Santa Cruz), and mouse anti-BirA (NBP2-59939; Novus Biologicals). Secondary antibodies used were goat Alexa Fluoro488-conjugated anti-mouse IgG (#A11001; Thermo Fisher Scientific), donkey Alexa Fluoro 647-conjugated anti-goat IgG (#A11001; Thermo Fisher Scientific), and goat Alexa Fluoro488-conjugated anti-rabbit IgG (#A11001; Thermo Fisher Scientific). Biotin was immunostained using Alexa Fluoro 647-conjugated streptavidin (#S21374; Thermo Fisher Scientific).

### Flow cytometry analysis

Cells were dissociated using Accutase (Merck), and incubated with mouse anti-EGFR antibody (#GR01; Merck Millipore) or mouse IgG control in 0.1% BSA/PBS for 30 min on ice. The cells were washed three times with 0.1% BSA/PBS, and incubated with Alexa488-conjugated anti-mouse goat IgG in 0.1% BSA/PBS for 30 min on ice. The cells were washed three times with 0.1% BSA/PBS, and resuspended in 0.1% BSA/PBS with DAPI (Thermo Fisher Scientific). The fluorescent signal was measured using LSRFortessa™ Cell Analyzer (BD Biosciences).

### Sample preparation for multilayered proteomics analysis (6 ligands analysis) Cell lysates

Clone T1 cells (4.0 × 10^6^ cells) were seeded in 15 cm dishes and cultured for 3 days. The medium was replaced by starvation medium. After 6 h starvation, the cells were stimulated with the indicated concentration of ligand samples for indicated time at 37°C. For washout method, medium was removed 20 min after the stimulation with ligands, washed by pre-warmed starvation medium twice, and replenished with pre-warmed starvation medium. Biotin is added 10 min before the harvest at the final concentration of 0.5 mM. After the stimulation, the cells were washed with ice-cold PBS and lysed using modified RIPA buffer supplemented with protease inhibitor cocktail. After the incubation on ice for more than 10 min, the lysates were scraped and centrifuged at 12,000 × *g* for 20 min at 4°C, and the supernatants were collected. The samples were divided into three groups depending on the ligands (EGF/TGFa, BTC/AREG, and HBEGF/EREG), and the lysates were prepared for one group on one day (called “experimental days’’ hereafter). The protein concentration of the cell lysates was measured using a BCA assay kit.

### Interaction network analysis samples

500 µg of proteins in modified RIPA buffer was incubated with 5 mM TCEP and 10 mM 2-Chloroacetamide (CAA) for 15 min at ambient temperature. The lysates were incubated with 100 µL of lysine-modified streptavidin magnetic beads for 2h, followed by the subsequent wash with 100 mM Tris-HCl (pH 8.5) and 0.1% Sodium dodecyl sulfate (SDS), 1 M KCl, 50 mM Tris-HCl (pH 8.5) and 3 M Urea, 50 mM Tris-HCl (pH 8.5) and 6 M Urea, and 70% ACN. The beads are eluted in 50 mM TEAB (pH 8.0) containing 100 ng of Lys-C and incubated overnight at 25°C. After the overnight digestion, the beads are removed and the supernatants were incubated with 100 ng of trypsin for 3h at 37°C. The solution was acidified with 10% formic acid (FA) and loaded on to Evotips for interaction network analysis.

### Phopshoproteomics samples

500 µg of proteins was incubated with 5 mM TCEP and 10 mM CAA for 15 min at ambient temperature, followed by the digestion using PAC method^2^ with elution in 50 mM TEAB (pH 8.0) containing 0.5 µg of Lys-C and 1 µg of Trypsin. After the digestion overnight at 37°C, the beads were removed, and the solution was acidified with 10% FA, loaded to Sep-Pak tC18 96-well µElution Plate for desalting, and eluted with 40% ACN and 60% ACN subsequently. The solutions were adjusted to 80% ACN/5% TFA/1 M glycolic acid (GA) and incubated with 20 µL of TiO^2^ beads. The beads were subsequently washed with 80% ACN/5% TFA/1 M GA, 80% ACN/1% TFA, and 10% ACN/0.2% TFA and eluted with 1% NH3 solution. The eluents were acidified with 10% TFA, filtered, and loaded on to Evotips for phosphoproteomics analysis.

### Proteomics samples

100 µg of proteins was incubated with 5 mM TCEP and 10 mM CAA for 15 min at ambient temperature, followed by the digestion using PAC method with elution in 50 mM TEAB (pH 8.0) containing 0.5 µg of Lys-C and 1 µg of Trypsin. After the digestion overnight at 37°C, the beads were removed, and the solution was acidified with 10% FA. The peptide concentrations were measured using NanoDrop and 750 ng of peptides were loaded onto Evotips.

### Sample preparation for multilayered proteomics analysis (chimeric ligands analysis)

Clone T1 cells (2.3 × 10^6^ cells) were seeded in 15 cm dishes and cultured for 2 days. The medium was replaced by starvation medium. After overnight starvation, the cells were stimulated, harvested, and processed to the following proteomics analyses by the same procedure as the “Sample preparation for multilayered proteomics analysis of 6 ligands“ section above.

### Sample preparation for interaction network analysis (EGFR tyrosine-mutant analysis)

HEK293T cells (4.4 × 10^5^ cells) were seeded in 6 cm dishes and transfected with 2.2 µg of the pCMVd2-EGFR-TurboID vectors using Lipofectamine 3000 the next day. The day after the transfection, the medium was replaced by starvation medium and cultured overnight. After the starvation, the cells were stimulated and harvested by the same procedure as the “Cell lysates“ section above. The lysates of the condition “Y1173F, EGF 20 min” were prepared on a different day from other lysates together with additional control samples. 200 µg of proteins were used for interaction network analysis using the same protocol as “Interaction network analysis samples”.

### LC-MS/MS and MS data analysis

Samples for all the proteomics analysis were analyzed by coupling the Evosep One system and Orbitrap Exploris 480 MS (Thermo Fisher Scientific) in data-independent acquisition (DIA) mode. Peptides were eluted in either an in-house packed 15 cm, 150 mm i.d. capillary column with 1.9 mm Reprosil-Pur C18 beads (Dr. Maisch) or a performance column (EV1137, EvoSep), and 30 samples per day (proteomics) or 60 samples per day (phosphoproteomics) were analyzed using a preprogrammed gradient. Temperature of the in-house columns was set to 60°C using an integrated column oven (PRSO-V1, Sonation), and that of the commercial column was to 40°C. The spray voltage was set to 2 kV, funnel RF level to 40, and capillary temperature to 275°C. Full MS resolution was set to 120,000 at m/z 200, full MS AGC target was set to 300% and IT to 45 ms. The mass range was set to 350-1400. AGC target for fragment spectra set to 100%; 49 windows with 13.7 m/z scans from 361 to 1033 m/z with 1 Da overlap. Resolution was set to 15,000, IT to 22 ms, and normalized collision energy was 27%; for phosphoproteome analysis using DIA, 16 windows of 39.5 m/z scans from 472 to 1143 m/z with 1 m/z overlap were used. Resolution was set to 45,000 and IT to 86 ms. Normalized collision energy was set to 27%. All data were acquired in profile mode using positive polarity.

### Bioinformatics of interaction network analysis (6 ligands analysis)

Proteins that were identified less than 66% of all the conditions were filtered out, and the dataset was log2-transformed and normalized using the “normalizecyclicloess” function of the “limma” package. The missing values of the dataset were categorize to partially observed values (POV) and “missing in the entire condition (MEV), and imputed using the “wrapper.impute.slsa” and “wrapper.impute.detQuant” functions of “DAPAR” package, respectively Proteins containing missing values that could not be imputed were filtered out after the imputation. Batch correction was performed by Combat using the experimental days and replicates as batch groups. Volcano plots were generated using Perseus software^3^ with the settings of FDR < 0.05 and s0 value = 0.1. “EGFR signaling” proteins in the volcano plots represent the proteins assigned by Perseus software in “ErbB signaling pathway” and “Endocytosis” in KEGG terms. The proteins were further filtered based on ANOVA significance (FDR < 0.01). Median values from 5 replicates are used as representative values for each protein in each condition, and PCA was performed in R using “prcomp” and “autoplot” function of the “ggfortify” package. Heatmap with hierarchical clustering was created in R using the “heatmap3” package with default settings. For each protein, those with a difference more than or equal to 1.5-fold between the maximum and minimum intensity out of all conditions were extracted, and at each time point, the intensity was averaged over the three ligands in each ligand category (recycling- and degradation-inducing ligands). Fuzzy c-means clustering was performed in R using the “Mfuzz” package^4^ with “Normal SD Based” standardization and the settings of “value of m = 1.5” and “number of clusters = 8”. The lists of proteins that are categorized into each cluster were used as inputs for GO and KEGG pathway enrichment analysis using DAVID^5^ with default background. “GOTERM_CC_ALL”, “GOTERM_BP_ALL”, “GOTERM_MF_ALL”, and “KEGG_PATHWAY” were selected as annotations. Enriched terms were filtered based on Bonferroni-adjusted p-value (p < 0.01), and the terms that show the significant enrichment in at least one cluster were plotted.

### Bioinformatics of phosphoproteomics (6 ligands analysis)

Phosphopeptides with a site-localization probability of at least 0.75 were included in the following analyses. Peptides that were identified less than 70% of all the conditions were filtered out, log2-transformed, normalization, and imputation was performed in the same manner as for interaction network analysis. Batch correction was performed by Combat using the experimental days as batch groups. Volcano plots were generated using Perseus software with the setting of FDR < 0.05 and s0 value = 0.1. “EGFR signaling” proteins in the volcano plots represent the proteins assigned in “ErbB signaling pathway” in KEGG terms by Perseus software. The proteins were further filtered based on ANOVA significance (FDR < 0.01). Median values from 5 replicates are used as representative values for each condition, and PCA was performed in the same manner as for interaction network analysis. Heatmap with hierarchical clustering was created in R using the “heatmap3” package with default settings For each phosphopeptide at each time point, the intensity was averaged over the three ligands in each ligand category. Fuzzy c-means clustering was performed in R using the “Mfuzz’’ package with “Normal SD Based” standardization and the settings of “value of m = 1.5” and “number of clusters = 10”. The sequences of 15 amino acids surrounding the phosphorylated sites were used for enriched phosphosite motif analysis using iceLogo^6^ with default settings. A list of the amino acid sequences from all the phosphopeptides before the filtration based on ANOVA significance was used as a background dataset for the motif analysis.

### Bioinformatics of proteomics (6 ligands analysis)

Proteins that are identified less than 70% of all the conditions were filtered out, log2-transformed, normalization, and imputation was performed in the same manner as above. Batch correction was performed by Combat using the experimental days and replicates as batch groups.

### Bioinformatics (Chimeric ligands)

Phosphopeptides with a site-localization probability of at least 0.75 were included in the following analyses. Proteins or peptides that are identified less than 70% of all the conditions were filtered out, log2-transformed, normalization, and imputation and batch correction were performed in the same manner as above. Median values from 5 replicates are used as representative values for each protein in each condition, and PCA was performed in R using “prcomp” and “autoplot” function of the “ggfortify” package. Volcano plots were generated using Perseus software with the setting of FDR < 0.05 and s0 value = 0.1. “EGFR signaling” proteins in the volcano plots represent the proteins assigned in “ErbB signaling pathway” in KEGG terms by Perseus software.

### Bioinformatics of interaction network analysis (tyrosine mutants)

Some samples had problems that C18 materials dried out before the sample acquisition, so these samples were removed from the following data analysis. Proteins that were identified less than 60% of all the conditions except the non-transfected condition and the condition of “Y1173F, EGF 20 min” were filtered out. The dataset was log2-transformed and normalized using the “width adjustment” function in Perseus software. At this point, the values from the “Y1173F, EGF 20 min” samples were merged to the dataset by following procedure. Before merging, the values from the smaller dataset, consisting of non-stimulated condition and “Y1173F, EGF 20 min” condition, were standardized by subtraction so that non-stimulated conditions have the same median values as in the larger dataset. After obtaining the complete dataset, proteins were further filtered by the following three steps. (1) Proteins with a difference more than or equal to 2-fold and p-value less than or equal to 0.05 comparing 20 min and unstimulated conditions. (2) Proteins with a difference more than or equal to 1-fold comparing 5 min and unstimulated conditions. (3) Proteins with a difference more than or equal to 1-fold between 5 min and unstimulated conditions. The filtering steps 1 and 2 were conducted using data from WT and all the 1F mutants, and the step 3 were done using only WT data. Median values from the replicates are used as representative values for each protein in each condition.

